# Viroscope: plant viral diagnosis from NGS data using biologically-informed genome assembly coverage

**DOI:** 10.1101/2022.09.14.507814

**Authors:** Sandro L. Valenzuela, Tomás Norambuena, Verónica Morgante, Francisca García, Juan C. Jiménez, Carlos Núñez, Ignacia Fuentes, Bernardo Pollak

## Abstract

Next-generation sequencing (NGS) methods are transforming our capacity to detect pathogens and perform disease diagnosis. Although sequencing advances have enabled accessible and point-of-care NGS, data analysis pipelines have yet to provide robust tools for precise and certain diagnosis, particularly in cases of low sequencing coverage. Lack of standardized metrics and harmonized detection thresholds confound the problem further, impeding the adoption and implementation of these solutions in real-world applications. In this work, we tackle these issues and propose biologically-informed viral genome assembly coverage as a method to improve diagnostic certainty. We use the identification of viral replicases, an essential function of viral life cycles, to define genome coverage thresholds in which biological functions can be described. We validate the analysis pipeline, Viroscope, using field samples, synthetic and published datasets and demonstrate that it provides sensitive and specific viral detection. Furthermore, we developed Viroscope.io a web-service to provide on-demand NGS data viral diagnosis to facilitate adoption and implementation by phytosanitary agencies to enable precise viral diagnosis.

## INTRODUCTION

Plant viruses are among the most important pathogens for agriculture, causing losses amounting up to more than $30 billion annually (Rao and Reddy, 2020; Rodríguez-Verástegui et al., 2022). Facile and accurate detection of plant viruses is essential to avoid propagation of these pathogens due to increasing global plant trade practices. Although methods such as next generation sequencing (NGS) can enable unbiased detection, these techniques are still in the process of being implemented. In addition, the standardization of analysis pipelines to perform plant virus diagnosis is pending, particularly in cases of low virus abundance (Jones et al., 2017; Massart et al., 2017; Massart et al., 2019; Mehetre et al., 2021).

Plant viral diseases are of considerable concern for farmers, researchers, and policy-makers since they are capable of decimating food production and even eradicating whole species (Legg et al., 2000; Gonsalves et al., 2008; Moreno et al., 2008). Viruses can have variable effects on the plant’s physiology, from the slight decline in productivity and quality of products to high levels of lethality. The latter was the case for the Citrus tristeza virus (CTV), which is estimated to have killed over 100 million plants over several countries worldwide (Jones R., 2021). Species such as sweet cherry (*Prunus avium*) have been traditionally multiplied using clonal propagation leading to accumulation of a large number of viruses. This is of particular concern since these pathogens may be latent or may cause detectable symptoms in susceptible rootstocks and/or scions (Umer et al., 2019). In addition, sweet cherry trees can have heterogeneous virus titrations during different seasons (Umer et al., 2019; Rodríguez-Verástegui et al., 2022).

Plant import and export practices demand strict phytosanitary controls to avoid propagation of pathogens between countries. In some cases, quarantines for up to several years are required before clearing plant material for import, creating a barrier to expedite transfer of newly developed varieties with improved traits. During quarantine, plants are monitored for evidence of viral symptoms as well as being repeatedly and directly tested for disease using molecular diagnostic assays (Jones R., 2009; Massart et al., 2017).

The two most successfully established plant virus detection methods are enzyme-linked immunosorbent assay (ELISA) and real-time polymerase chain reaction (PCR, or quantitative qPCR). ELISA detects structural protein motifs of the virus and can have a broad capability to detect variants, but exhibits limited sensitivity of detection for cases of low virus abundance. On the other hand, PCR-based assays (i.e.: PCR, RT-PCR, qPCR) are considered the gold standard for detection of a virus presence, however it requires *a priori* knowledge of virus target sequences. Thus, PCR-based methods are biased, depending on the availability of the sequences to design the analysis and can fail to detect virus variants. In spite of the advantages or disadvantages of these two methods, both techniques show high levels of reproducibility, capability for automation and are relatively low-cost for industry standards (Boonham et al., 2014; Chauhan et al. 2019,).

Methods such as next-generation sequencing (NGS) can help to address food security from increasing threats of viral disease outbreak (Massart et al., 2017, Mehetre et al., 2021). NGS enables unbiased and hypothesis-free testing of plant samples, and is becoming increasingly cost effective, with price per base pair sequenced dropping dramatically over the past decade. Viruses detection by NGS comprises: (1) nucleic acid extraction from the plant material, (2) library preparation (enriching virus sequences or depleting host sequences such as ribosomal RNA), (3) high-throughput sequencing, (4) raw data quality control and removal of poor quality reads and adaptor sequences, (5) removal of host reads, (6) mapping to a virus database, and/or *de novo* assembly of reads, (7) read counting and/or coverage calculation, (8) identification of present viruses using read and coverage cutoffs (Villamor et al., 2019). Specifically, the bioinformatic identification pipeline (involving aforementioned steps 4 through 8) is essential for accurate diagnosis. The potential of NGS has been recognised by several phytosanitary agencies where scientific committees and workshops have been held to address how to implement these techniques and the challenges involved. Furthermore, international organizations towards plant diseases protection are looking to improve the availability of diagnostic tools and are currently revising their diagnostic standards (Adams et al., 2018; Jones and Naidú, 2021).

The success of a NGS-based virus diagnosis is highly dependent on proper computing infrastructure and bioinformatics expertise (Jones et al., 2017; Umer et al., 2019; Kutnjak et al., 2021). Also, substantial virology knowledge is required to suitably interpret the results (Gaafar et al., 2021). Although the vast majority of virus diagnostic tools have been focused on human clinical samples –*VirusFinder* and *VERSE* (Wang et at., 2013; 2015), *VIP* (Li et al., 2016), *VirusSeeker* (Zhao et al., 2017)– some efforts have been focused specifically on plants –*VirFind* (Ho and Tzanetakis, 2014), *VSD toolkit* (Barrero et al., 2017), *Virtool* (Rott et al., 2017) and *PVDP* (Gutierrez et al., 2021). However, bioinformatic pipelines have presented challenges for standardization and incongruences in frequently used metrics such as read counts, genome coverage or a combination of both criteria are common (Visser et al., 2016, Rott et al., 2017, Malapi-Wight et al., 2021, Soltani et al., 2021, Hanafi et al. 2022). Also, threshold harmonization is required to establish virus detection using NGS (Ruiz-García, et al., 2021), particularly in cases with low sequencing coverage.

Low sequencing coverage presents a complex diagnostic challenge for virus detection. It may be due to a number of causes, such as low viral titre, insufficient depth of sequencing, sample cross-contamination, remnants of a past infection or even a latent phase of a virus. Most plant viruses have an RNA genome that adopts a basic replication mechanism consisting in the RdRp enzyme (RNA-dependent RNA Polymerase) as the responsible for transcription and replication (Hull, 2014). In other words, the identification of such an essential biological functionality in viruses may aid in interpreting cases of low sequencing coverage, as it implies capacity for propagation and thus, potential infectivity.

Here, we present Viroscope, a diagnostic pipeline that improves virus detection accuracy by using biologically-informed viral genome assembly coverage (VGAC). We introduce the identification of replicases to evaluate how different VGAC threshold levels correlate with functional aspects of virus biology. In addition, we evaluate the performance of Viroscope with field samples of sweet cherry, simulated datasets and external datasets, demonstrating that VGAC is a robust measure for virus detection using total RNA NGS data using Illumina and Nanopore sequencing. Finally, we have implemented the pipeline in the form of a web application called *Viroscope.io* (https://www.viroscope.io) to enable cloud-based NGS data virus diagnosis.

## MATERIALS AND METHODS

### Collection of field samples and nucleic acid extraction

Sweet cherries are one of the major stone fruits cultivars in south-central Chile. For this study, a sweet cherry production farm located in the O’Higgins region was selected. The plant material was randomly collected from four different >5 years old *P. avium* specimens. In this field, elite varieties ‘Lapins’ and ‘Santina’ were the most common cultivars. Hence, each sample was called L1, L2 and S1 and S2, respectively. In order to evaluate differences in plant virus diagnosis due to season conditions (factors such as changes in temperature, light, and/or plant nutrition), the same plants were sampled at two different principal growth stages as previously described for *Prunus sp*.: Stage 3 or shoot development (SD) and Stage 9 or senescence (SS, beginning of dormancy) (Fadón et.al., 2015). Each growth stage corresponds to the Spring and the end of Summer season, respectively. A total of eight leaves (four apical and fourequal-sized middle-aged) from the canopy of an individual tree were sampled and placed in RNAlater solution (Invitrogen). Then, the samples were transported to the laboratory in refrigerated containers and stored at −80°C until their use.

For total RNA extraction, leaves in RNAlater were pooled and ground in a liquid nitrogen cooled ceramic mortar, and 100 mg of ground sample was processed using the Spectrum™ plant total RNA kit (Sigma-Aldrich, St. Louis, MO, USA), according to the manufacturer’s instructions. Total RNA concentration and quality was assessed using an optical microplate reader (BioTek Synergy H1, Santa Clara, California, USA), through fluorometry (Promega Quantus fluorometer, Madison, Wisconsin, USA) and integrity was verified through agarose gel electrophoresis (Rio, 2015). Purified extracts were stored at −80°C until further processing.

### RNA-sequencing of field samples

For RNA-sequencing, 2 μg of total RNA from each of the field samples was sent to Novogene Corporation Inc. (USA) (samples SD-L1, SD-L2, SD-S1 and SD-S2) and Macrogen Co., Ltd (Korea) (samples SS-L1, SS-L2, SS-S1 and SS-S2). Ribosomal RNA depletion was performed using Ribo-Zero Plant (Illumina, USA), and library construction was executed using NEBNext Ultra RNA Library Prep kit (New England Biolabs, USA), and sequenced on a NovaSeq6000, using a 150 bp paired-end cycle.

### NGS datasets from field samples and processing

Raw reads from field samples (SD/SS-L1, SD/SS-L2, SD/SS-S1, SD/SS-S2; Supplementary Table S1) were filtered using *fastp* (Chen et al., 2018) keeping reads with an average quality greater or equal to 20. Single reads, reads shorter than 50 bp and reads containing “N” nucleotides were discarded, and adapters were automatically trimmed.

In order to assay the dependency of the prediction on the size of data, sequencing data of field samples were subjected to a *jackknife* process. Reads were randomly selected from the original data to obtain sets of different depths of sequencing, namely 100K, 500K, 1M, 5M, 10M and 15M reads. Each selection was repeated 10 times. Subsampling read repetitions as well as sequence manipulation were performed using the *seqtk* subsampling routine (using the repetition number as a seed for paired reads) and *seqkit* (Shen et al., 2016).

### NGS datasets from simulation and subsampling

Artificial NGS datasets of 20M paired-end reads of 150 bp (Supplementary Table S1) were generated using the software *ART* (Huang et al., 2012) and *seqkit* (Shen et al., 2016). Two of these datasets were intended to simulate actual field samples containing the 11 viruses of the Pavium panel-I (see section *Read assignment and viral panels* below). In the first dataset (Synab dataset), virus abundance was based on experimental data (average distribution as seen in real sequencing data of samples SD-L1, SD-L2, SD-S1 and SD-S2), whereas the second dataset (Synhom dataset) assumed an even distribution of virus read abundance (Supplementary Table S2 and Supplementary Table S3). Both sets were then subjected to subsampling to get different depths of sequencing (100K, 500K, 1M, 5M, 10M and 15M reads) in a *jackknife* process (10 repetitions).

Another dataset of 20M paired-end reads (150 bp) was generated (Mut dataset), where all the viruses (from Pavium panel-I) were randomly mutated at different rates (5, 10, 15, 20, 25, 30%). In this case, a *jackknife* process with subsampling of 10M reads each was done for 10 repetitions (using repetition number as seed for subsampling). *Mutation-Simulator* (version 2.0.3) (Kühl et al., 2021) was used to simulate the mutations on the viruses.

Two additional datasets were built to further investigate the relationship between virus abundance and depth of sequencing. In this case, datasets of 20M total paired-end reads (150 bp) containing at most 3% of viral reads (600K reads) from the viruses with the shortest (Cherry Virus A, CVA) and the longest (Little Cherry Virus 1, LChV-1) genomes from the Pavium panel-I were surveyed (Cva and Lchv1 datatsets). Subsets containing 0.05, 0.1, 0.5, 1, 5, 10% of such 3% of representation of the viral reads defined above were generated. For example, a 20M total read set with 10% of 3% viral reads contains 0.1 × 0.03 × 20M = 60K viral reads, while at 0.5% of 3% viral abundance it contains 3K reads. The remaining reads consisted of background sequences from plant, human, bacteria, and random sequences in the same proportion as listed in Supplementary Table S2. To compensate for the decreasing number of reads, the difference was replaced by bacterial reads. Each of these sets was in turn subsampled in a *jackknife* process to obtain 10 subsets at different depths of sequencing (100K, 500K, 1M, 2M, 3M, 4M, 5M, 6M, 7M, 8M, 9M, and 10M random reads).

### NGS datasets from published reports

Several external datasets containing NGS data were surveyed and tested (Supplementary Table S1). A total of 32 datasets were tested: ten datasets are part of a challenge for identifying viruses in NGS data under different conditions (Tamisier et al., 2021; single-end (SE) or paired-end (PE) semi-artificial short reads); one dataset is part of an analysis using a plant transcriptome to identify viruses (Jo et al., 2016; RNAseq, SE reads); one dataset comes from the study of the pepper virome (Jo et al., 2017; RNAseq, PE reads); one dataset is part of a study of small RNAs produced by Dicer-like enzymes as a defense strategy of a plant when infected by a virus (Barrero et al., 2017; sRNAseq, SE reads); 14 datasets come from a report describing the use of Oxford Nanopore’s MinION to detect and genotype potato viruses (Della-Bartola et al., 2020; RNAseq, ONT reads); one dataset intended to report the genome sequence of a virus based on Oxford Nanopore (Leiva et al., 2020; DNA, ONT reads); three datasets accounting for the identification of genomes of viruses affecting crops in sub-Saharan Africa (Boykin et al., 2018; DNA, ONT reads); and one dataset from a study that demonstrates the use of MinION sequencing to detect and characterize viruses infecting water yam plants (Filloux et al., 2018; RNAseq, ONT reads). All the datasets contain a sum of 46 different viruses to be detected and 62 cases (a “case” is defined as a “virus to be detected in a dataset”, for example, there are two cases when virus A is present in dataset 1 and 2, or when virus A and virus B are present in dataset 1), which were divided into three groups (some of the viruses are included in more than one dataset): Viromock datasets V1-V10 (18 viruses; 15 cases), SmallRNA datasets R1-R3 (21 viruses and one viroid; 22 cases), and Nanopore datasets N1-N19 (10 viruses; 25 cases). All the viruses that must be detected in each group as well as additional details of these datasets are listed in Supplementary Table S4.

### Read assignment and viral panels

For read assignment (taxonomic classification of reads according to a panel of virus genomes), two widely used software for metagenome exploration were tested, namely *Kraken2* (Wood et al., 2019) and *Centrifuge* (Kim et al., 2016), which are the fastest and more sensible algorithms according to a benchmark previously published (Miossec et al., 2020). *Minimap2*, a classical algorithm used to compare local read alignment (Li H., 2018) was also tested in the read assignment process. All software was run with default parameters, except *Kraken2* whose database was built by lowering the default parameter k (*kmer* length) from 35 to 31 to increase its sensitivity.

For all the dataset groups (Supplementary Table S1), *ad hoc* viral panels were built in order to set the proper databases required by each software. In the case of both the field samples (SD/SS-L1, SD/SS-L2, SD/SS-S1, SD/SS-S2) and simulated datasets (Synab, Synhom, Mut, Cva, Lchv1), the analysis was carried out using 11 viruses which affect *Prunus sp*., some of which were previously reported in Chile: Apple chlorotic leaf spot virus (ACLSV), Apple mosaic virus (ApMV), Cherry green ring mottle virus (CGRMV), Cherry necrotic rusty mottle virus (CNRMV), Cherry virus A (CVA), Little cherry virus 1 (LChV-1), Plum bark necrosis stem pitting-associated virus (PBNSPaV), Prune dwarf virus (PDV), Prunus necrotic ringspot virus (PNRSV) and Plum pox virus (PPV) (Fiore et al., 2016), and absent such as Little cherry virus 2 (LChV-2). This set of viruses was called “Pavium panel-I” and the respective database included reference sequences from NCBI (Supplementary Table S3). Additionally, an extended database of this panel was built by incorporating the different isolates of the 11 viruses. A total of 1,011 sequences were obtained from NCBI (including the original 11 reference sequences) using all sequences under defined taxID and keeping only complete genomes, which were clustered with *CD-HIT* (Fu et al., 2012). Using the requisite of 90% sequence identity, 139 clusters were obtained, whose representative sequences became the Pavium panel-II (Supplementary Table S5). Although a lower sequence identity can account for the same virus species, the rationale of this requirement is just to identify virus species considering the possible sequence differences, which is not affordable when using only reference sequences. The use of 90% of sequence identity comes from the mutation simulation analysis (see Results), which is a proper trade-off between incorporating more virus isolates into the panel and not including all the sequences.

Regarding the external published datasets, viral panels and the respective reference databases were built for each group according to the viruses that must be detected in each of them (Supplementary Table S4). Therefore, the panels Viromock (18 viruses: BPEV, BSV, BYDV, CMV, CTV, CVEV, EMDV, GRBaV, GRLaV2, GRSPaV, GRVFV, LChV-1, PBNSPaV, PepMV, PFBV, PiVB, PVY, and TSWV), SmallRNA (21 viruses: AGCaV, ALPV, ASGV, ASPV, BPEV, CLCuV, CYVMV, GalLV, GFkV, GLRaV-3, GRSPaV, GVB, PepLCBV, PepLCVB, PeSV, PrVT, ToLCBDB, ToLCGV, ToLCJoV, ToLCRnV, and TVCV) and Nanopore (10 viruses: CMV, DBV, EACMV, PLRV, PVS, PVX, PVY, SLCMV, YCNMV, and YMMV) were created. In the case of the Viromock datasets, the viruses present in the datasets V11-V18 were included in the panel, but the datasets were not surveyed since they were generated for identification of viral isolates and did not contain background sequences.

### Viral genome assembly coverage (VGAC)

Reads assigned to a viral reference sequence were *de novo* assembled. *SPAdes* (Bankevich et al., 2012) was used to perform the assembly on Illumina reads (in ‘--sc’ mode and with ‘--careful’ option), whereas *Canu* (Koren et al., 2017) was used to assemble Oxford Nanopore reads. In either case, contigs obtained in the assembly process were then remapped to the respective reference genome. In this case, the mapping was performed using *Minimap2* (with ‘-map-ont’ option to map contigs), and the genome coverage (percentage of the reference genome that is covered by the assembled contigs) was calculated by *SAMtools/BCFtools* (Danecek et al., 2021), with the following commands:

~~~
$ minimap2 -ax map-ont reference_genome.fasta contigs.fastq -o aligned_contigs.sam
$ samtools view -bh -o aligned_contigs.bam aligned_contigs.sam
$ samtools coverage aligned_contigs.bam
~~~

And the VGAC was calculated as:

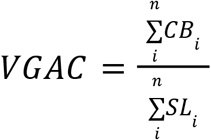

where, *CB_i_* is the number of covered bases with depth ≥ 1 for the segment *i* of the virus, and *SL_i_* is the length of the segment *i* of the virus (most of the viruses contain only one genome segment, so *n = 1* in these cases). Thus, the VGAC is a value that ranges between 0 and 1 (and can also be expressed as percentage).

### Replicase identification

Consensus regions were determined from mapped contigs onto the reference genome of the virus using *SAMtools/BCFtools*. These consensus regions were calculated as follows:

~~~
$ samtools mpileup -uf reference_genome.fasta -o mpile.vcf aligned_contigs.bam
$ bcftools call -c --ploidy 1 -o call_mpile.vcf mpile.vcf
$ vcfutils.pl vcf2fq call_mpile.vcf > consensus.fastq
~~~

Each consensus region was compared against a repository of protein sequences related to a virus replication (e.g. replicase or polymerase). This repository was built from *RVDB-prot* (version 23.0, 2021-12; Bigot et al., 2020) and contains 17,708 records. To build this repository, *RVDB-prot* was filtered using terms accounting for replicase or polymerase activity. The terms searched for were: ”replicase”, “RNA dependent RNA polymerase” (RdRp), “RdRp”, “RNA dependent DNA polymerase” (RdDp), “RdDp”, and “polymerase”, which allowed for recovering 13,576 sequences. In addition, the term “reverse transcriptase” (which is a synonym of “RdDp”) was searched, producing 59,525 records. In this case, records belonging to “homo”, “human”, “hepatitis”, and “hiv” were filtered out, remaining 4,114 sequences, which were incorporated into our repository. In the course of evaluating the different panels, some of the virus sequences did not account for the presence of a coding sequence related to replicase or polymerase activities. However, all these cases were manually inspected and found to encode in fact a protein either with a name containing none of the terms searched above, being absent in *RVDB-prot* or being part of a polyprotein. Finally, all of them were incorporated into our viral replicase protein repository.

The comparison of the consensus regions was performed using *Diamond* (Buchfink et al., 2021) with its *blastx* module. The resulting hits were then filtered by similarity (80 or 90%) and length of alignment (80 or 90%). At the 80/80 schema, a replicase was said to be identified in the respective consensus region if it contained 80% similarity and 80% sequence alignment (90/90 represents a more stringent schema).

### Overall performance on external datasets

In order to estimate the performance of the different stages of the pipeline on the external datasets, several measures were calculated: sensitivity, specificity, precision, accuracy, and false discovery rate (FDR). These measures were determined as follows:

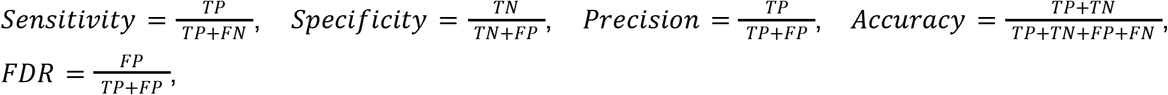

where *TP* (true positives) is the number of cases that were detected that must be detected, *FP* (false positives) is the number of cases that were detected that should have been not detected. In this sense, for negatives, *FN* (false negatives) corresponds to the number of cases that were not detected that should have been detected, and *TN* (true negatives) is the number of cases that were not detected that in fact were not present. Since datasets are reported to have only the viruses to be detected, *FP* and *TN* values were estimated from the rest of viruses of the respective panel which should account for misassigned reads by at least one software. For example, in the Viromock datasets there are 15 actual cases to be detected, but the read assignment together yielded 51 additional cases detected, so a maximum of 51 *TN* cases were assumed to exist in these datasets. The measures listed above were determined for read assignment (the case is assumed to be a *TP* when existing at least one assigned read), and for detection of replicases (the case is assumed to be a *TP* either when matching the 80/80 or the 90/90 schema). In the case of VGAC, the measures were calculated for the thresholds 0.1 (the case is assumed to be a *TP* when VGAC ≥ 0.1), 0.2, 0.3, 0.4, 0.5, 0.6, 0.7, 0.8, 0.9, and 1.0. Finally, in order to estimate the similarity between measures for VGAC and the detection of replicases, the euclidean distance *d* between the measures for VGAC and for the schema 80/80 was determined as:

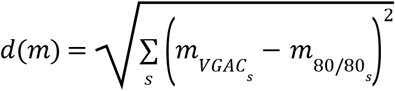

where *m* could be sensitivity, specificity, precision, accuracy, or FDR, and *s* runs on the three software (*Centrifuge, Kraken2* and *Minimap2*).

### Reverse transcription-polymerase chain reaction (RT-PCR) analysis

The presence of the viral pathogens in each individual sample collected at shoot development stage (SD-L1, SD-L2 and SD-S1 and SD-S2) and senescence stage (SS-L1, SS-L2 and SS-S1 and SS-S2) were confirmed by RT-PCR. The two-step RT-PCR for SD samples (Shoot Development) was performed by Laboratorio de Virología, Universidad de Chile (N. Fiore, personal communication, October 21, 2020), named here as “PCR external”. On the other hand, in the case of samples obtained during the senescence stage (SS-L1, SS-L2 and SS-S1 and SS-S2), the presence of the viral pathogens were confirmed by a two-step RT-PCR performed in this study. This method was optimized to detect the 11 viruses of the Pavium panel-I, and results were recorded as “PCR internal”. Primers of this panel were exclusively designed for this study based on a local viral sequences database and OligoPerfect™ designer software (ThermoFisher Scientific) (Supplementary Table S6). The phytoene desaturase 1 (*PDS1*) is a plant gene that exhibits constitutive expression and was used in our RT-PCR experiments as an internal control for RNA extraction and reverse transcription. In addition, the 11purified amplicons were used for positive controls to corroborate amplification of molecular targets of appropriate size (Supplementary Figure S6 E).

First-strand cDNA synthesis was performed using 70 ng of total RNA. The reverse transcription (RT) mix contained 200 units of recombinant Moloney Murine Leukemia Virus (MMLV) reverse transcriptase (Promega), 20 units of RNAsin (Promega), 1 mM dNTPs, and 1 uM of random hexamers (Promega). The reaction was performed at 20 uL final volume and was incubated at 37 °C for 60 min followed by enzyme inactivation at 70 °C for 5 min. The PCR mix (final volume of 25 uL) contained 1 uL of the cDNA, 1X (2,5 uL) GoTaq G2 Flexi Buffer (Promega), 0.15 uM of each primer (Supplementary Table S6), 3 mM MgCl2, 0.2 mM dNTPs, and 1.25 units of GoTaq G2 Flexi DNA polymerase (Promega). Cycling conditions for all primer pairs consisted of initial denaturation at 95 °C for 2 min followed by 35 cycles at 95 °C for 15 sec, 60 °C for 30 sec, 72 °C for 1 min and a final extension at 72 °C for 5 min. PCR products were analyzed by gel electrophoresis using 3% agarose in a 1X TBE buffer, and staining with 1:10.000 v/v SybrSafe (Invitrogen Life Technologies).

## RESULTS

### Overview of Viroscope

The Viroscope pipeline consists of two distinct steps for plant virus diagnosis based on NGS data (Figure 1A). First, a rigorous data analysis step encompassing: (1) read assignment, (2) *de novo* assembly of assigned reads, (3) reference mapping of assembled contigs, (4) genome coverage calculation of mapped contigs, (5) consensus calling, and (6) replicase identification in consensus sequences. In a second step, Viroscope detects pathogens by considering the VGAC values obtained by three read assignment algorithms and the identification of replicases. The validation of the pipeline was performed with three types of datasets: field samples from a sweet cherry farm, simulated datasets, and publicly available datasets (including different library preparation methods and sequencing technologies). The Viroscope results for field samples were also validated using RT-PCR methods to compare and study diagnostic sensitivity according to the different cutoff levels investigated. The Viroscope algorithm and the experimental design of this study are shown in Figure 1B.

**Figure 1.**
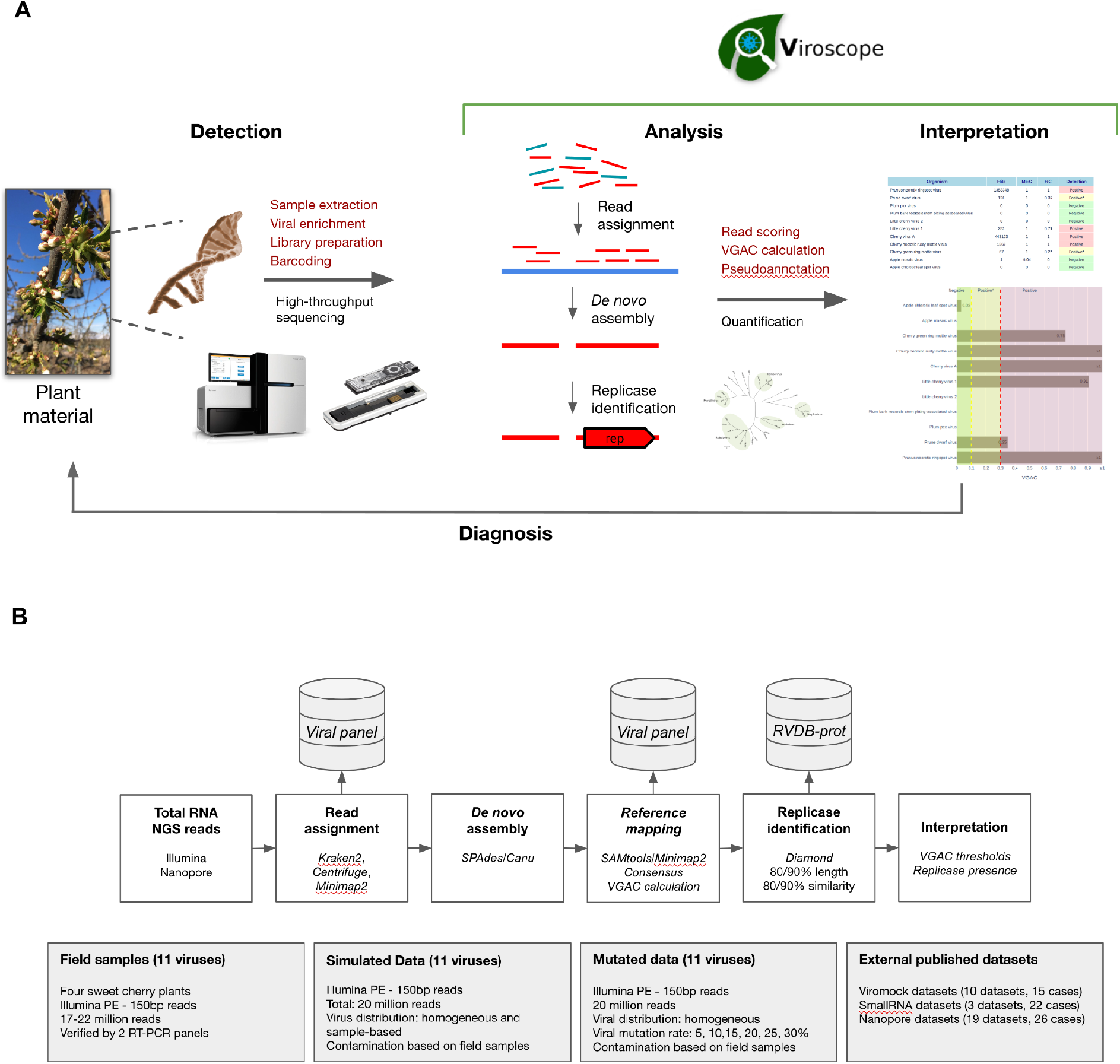
Overview of Viroscope, the data analysis pipeline and experimental validation strategy. **A**. Overview of Viroscope. Viroscope is a total RNA NGS data analysis pipeline that enables accurate viral detection by performing read assignment, *de novo* assembly with reference-based mapping and pseudo-annotation to obtain VGAC metrics and identification of viral replicases. These metrics inform the interpretation of viral presence from NGS reads to provide accurate diagnosis, contributing to the implementation of NGS for plant viral diagnosis in real-world applications. **B**. The Viroscope pipeline and experimental validation strategy. Viroscope performs read assignment from total RNA NGS (Illumina or Nanopore) reads using 3 read assignment software against a curated database of target viruses. Then, mapped reads are collected and used for de novo assembly either using *SPAdes* or *Canu* (for Illumina and Nanopore reads, respectively). Assembled contigs are used to perform reference-based mapping to obtain a consensus of mapped contigs to calculate VGAC. Then, the pipeline searches for the presence of replicases using Diamond using the RVDB-prot database. Finally these metrics are used for interpretation of viral presence according to specific cutoffs for diagnosis. Four experimental sets were used to validate the pipeline, sweet cherry field samples, a simulated dataset, a mutation dataset and external published datasets.

### Read assignment in field samples

In order to assess the viral abundance in sweet cherry samples, read assignment was performed using three different software, namely *Centrifuge, Kraken2* and *Minimap2*. The goal of this study was not to evaluate the different software, but to incorporate more than one perspective in the analysis since different algorithms could yield diverse results. Samples from four cherry plants specimens (SD-L1, SD-L2, SD-S1 and SD-S2) collected at the shoot development stage were sequenced using Illumina, yielding 17M-22M paired-end reads each (Supplementary Table S1), and read-subsampling was performed to evaluate the relationship between read assignment and depth of sequencing. The read assignment was carried out using a reference database called “Pavium panel I” comprising 11 viruses, namely ACLSV, ApMV, CGRMV, CNRMV, CVA, LChV-1, LChV-2, PBNSPaV, PDV, PNRSV and PPV (Fiore et al., 2016).

According to the read assignment process, the number of mapped reads increased in regard to the depth of sequencing in a linear fashion (Figure 2A and Supplementary Figure S1). Differences were observed according to the bioinformatic tool used: *Centrifuge* and *Kraken2* showed higher read assignment in relation to *Minimap2*. In addition, read assignment by only one software was observed for the case of CGRMV in sample L1, whose reads were assigned by *Centrifuge* but not by *Kraken2* nor by *Minimap2* (Figure 2A). Similar discrepancies were obtained in the cases of ACLSV for SD-L1 (reads assigned only by *Kraken2*), ApMV for SD-L1 (by *Centrifuge*), LChV-1 for SD-L1 (by *Kraken2*), LChV-1 for SD-L2 (by *Kraken2*), ACLSV for SD-S1 (by *Centrifuge*), ApMV for SD-S1 (by *Centrifuge*), and LChV-1 for SD-S1 (by *Kraken2*), although with a relatively low number of reads (Supplementary Figure S1).

**Figure 2.**
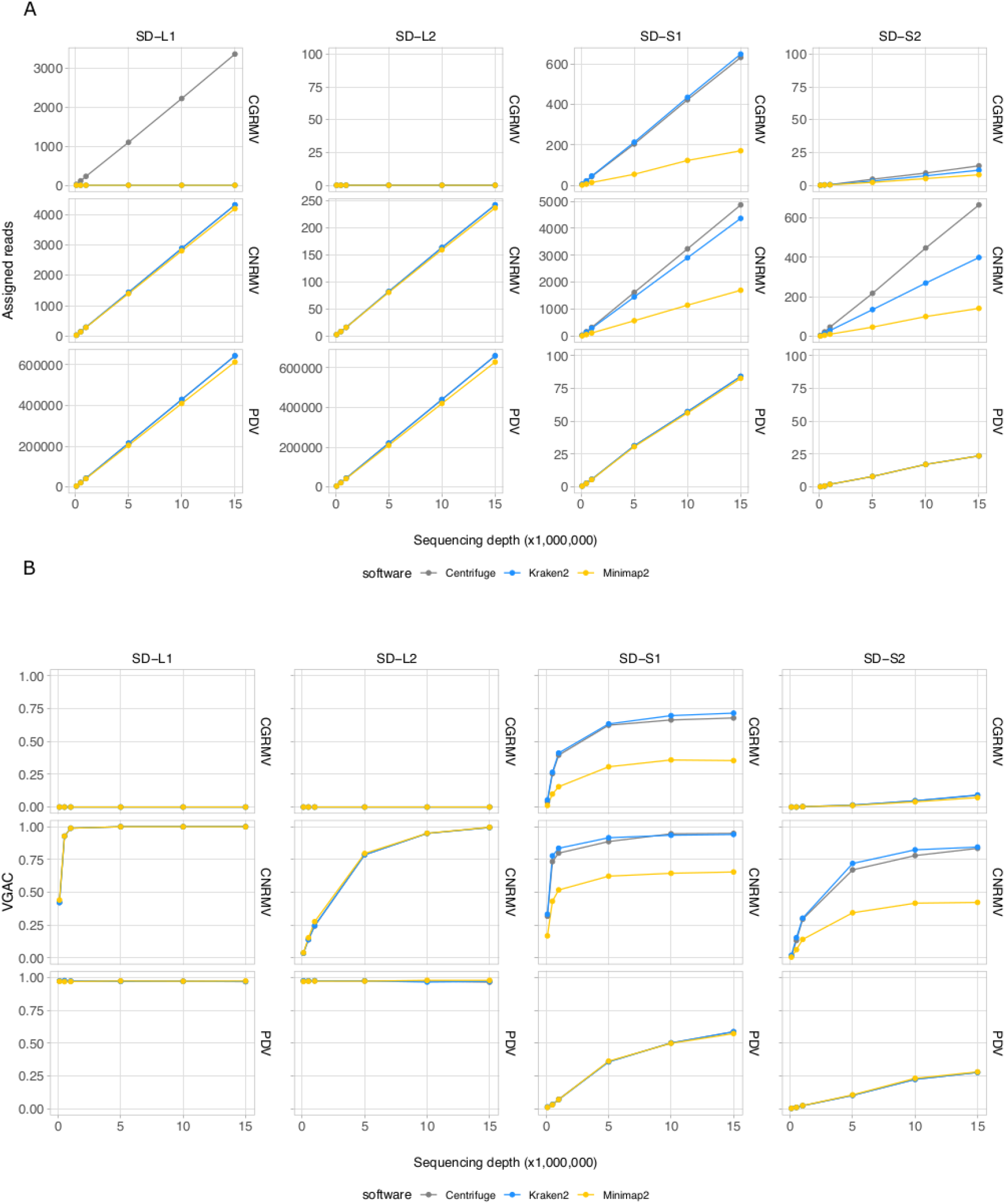
Read assignment and assembly coverage with NGS data from field samples. Ten subsets of randomly selected reads from sequencing data for field samples of cherry plants at the shoot development stage (SD-L1, SD-L2, SD-S1, and SD-S2) were built at different depths of sequencing. Three bioinformatic algorithms were tested, namely *Centrifuge, Kraken2*, and *Minimap2*. All the dots represent the average of 10 measures. VGAC was calculated according to Materials and Methods and with the reads assigned by the different algorithms at the respective depth of sequencing. **A**. Read assignment at different depths of sequencing (note the different scales of the ordinates) using the Pavium panel-I. **B**. VGAC obtained from the assembly of assigned reads (ranges from 0 to 1). Only the cases for CGRMV, CNRMV, and PDV are presented, but full versions containing all of the target viruses are depicted in Supplementary Figure S1 and Supplementary Figure S2. Average values (dots) as well as standard deviations are listed in Supplementary Table S11.

### Viral genome assembly coverage (VGAC) in field samples

Inspection of read assignment by the different software and their relation with assembly coverages at different depth of sequencing was carried out in order to evaluate and analyze detection issues. This was performed to study cases of low abundance of reads or possible misassignment. The detection of viruses was assessed through the VGAC, which accounts for a full or partial viral genome recovery using NGS data. As shown in Figure 2B, the virus detection was dependent on the sample and the abundance of viral sequences, for instance the cases CNRMV for SD-L1 (VGAC = 1.0 at 1M reads), CGRMV for SD-S1 (VGAC ≈ 0.72 at 15M reads), CNRMV for SD-S1 (VGAC ≈ 0.90 at 15M reads), PDV for SD-S1 (VGAC ≈ 0.60 at 15M reads), CNRMV for SD-L2 (VGAC = 1.0 at 15M reads), CGRMV for SD-S2 (VGAC ≈ 0.10 at 15M reads), and CNRMV for SD-S2 (VGAC ≈ 0.85 at 15M reads). In all the cases where only one software was able to assign reads and VGAC resulted to be ≈ 0 (e.g. CGRMV for SD-L1) (Supplementary Figure S2), the assigned reads could not assemble contigs, suggesting these were spurious or misassigned reads.

### Read assignment in simulated data

Depth of sequencing and the abundance of viral reads certainly influence the capability to perform the virus detection. In order to study the relationship between sequencing coverage and the VGAC, two synthetic NGS datasets with known viral composition were created for further analysis. The first dataset contained reads from fiveviruses based on the mean viral distribution of the field samples (Synab dataset) and the second one contained a homogeneous distribution of reads from Pavium panel-I, which are composed of 11 viruses (Synhom dataset). In order to mimic real samples, both datasets were built so that they contained background sequences, that is, they were contaminated with reads derived from human, plant, bacteria and random sequences. In both cases, the total quantity of viral reads was limited to 3% of total reads as shown in the distribution listed in Supplementary Table S2 and Supplementary Table S3, which in turn is based on the average empirical distribution. When a homogeneous distribution was used (*synhom* data in Figure 3), no differences in read assignment and VGAC amongst all viruses was observed (see Supplementary Figure S3 for more details). As expected, higher depth of sequencing resulted in higher read assignment consistent with what was seen in the field samples. These results were independent of both the virus (e.g. length or number of segments) and the assessed software, and no differences were observed regarding the VGAC values either. Moreover, complete viruses were assembled even at the lowest depth of sequencing (100K reads).

**Figure 3.**
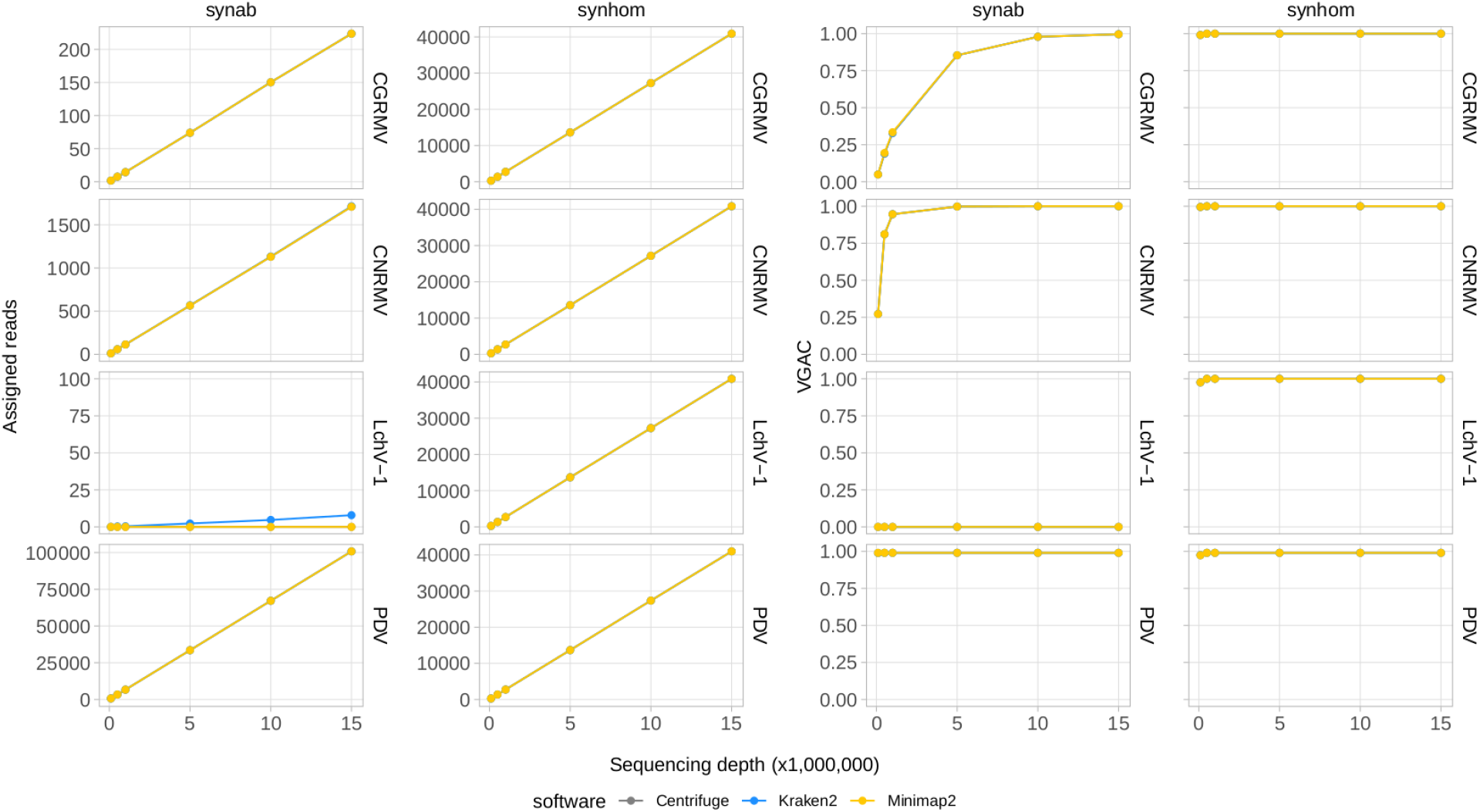
Read assignment and coverage in synthetic data. Synthetic NGS data were generated to simulate actual samples. The Pavium panel-I comprising 11 viral genomes was used to build samples containing 3% of viral reads from a 20 million paired-end reads. Samples were built with both a homogeneous distribution of viral reads (*synhom*) and based on an average distribution of actual samples (*synab*) (Supplementary Table S2 and Supplementary Table S3). In all cases, 10 subsets of randomly selected reads were built at different depths of sequencing. Assigned reads are shown in the two first column charts and VGAC is shown in the last two column charts. A full version containing all of the target viruses is depicted in Supplementary Figure S3. Average values (dots) as well as standard deviations are listed in Supplementary Table S11.

In the simulated data with empirical distribution (*synab* data in Figure 3), read assignment was proportional to the depth of sequencing and to viral abundance. Viruses used in the simulated data were CGRMV, CNRMV, CVA, PDV, and PNRSV (those with percentage > 0 in Supplementary Table S3), so reads assigned to the LChV-1 genome were considered as a misassignment. In fact, this virus showed no assembly (VGAC = 0) at each depth of sequencing, confirming such a misassignment (Figure 3). These results are consistent with the above observation that the VGAC was sensitive to both the depth of sequencing and the relative abundance of the viral reads in the field samples (e.g. the cases for CGRMV and CNRMV).

### Read assignment and VGAC in simulated mutation data

Further simulated datasets were generated so as to assess the tolerance of the different software to viral mutation rates. Subsets of 10M paired reads sampled from a set of 20M reads were used. At a 10% mutation rate, all the software were able to assign reads, and even at 20%, there was still a portion of an average ~2,900 out of 27,000 reads assigned to the viral genomes (Figure 4 and Supplementary Figure S4). Read assignment was not dependent on the viruses, which were homogeneously distributed in the simulated samples. Amongst the algorithms assessed, *Centrifuge* outperformed in the read assignment process in general, which was more evident at higher mutation rates. In this case, *Minimap2* resulted to be the least tolerant tool towards mutations. Although an increase in the mutation rate appeared to have more impact on the read assignment, at a 20% mutation rate the number of reads was still sufficient to assemble contigs at least with *Centrifuge* and *Kraken2* (VGAC ≈ 1).

**Figure 4.**
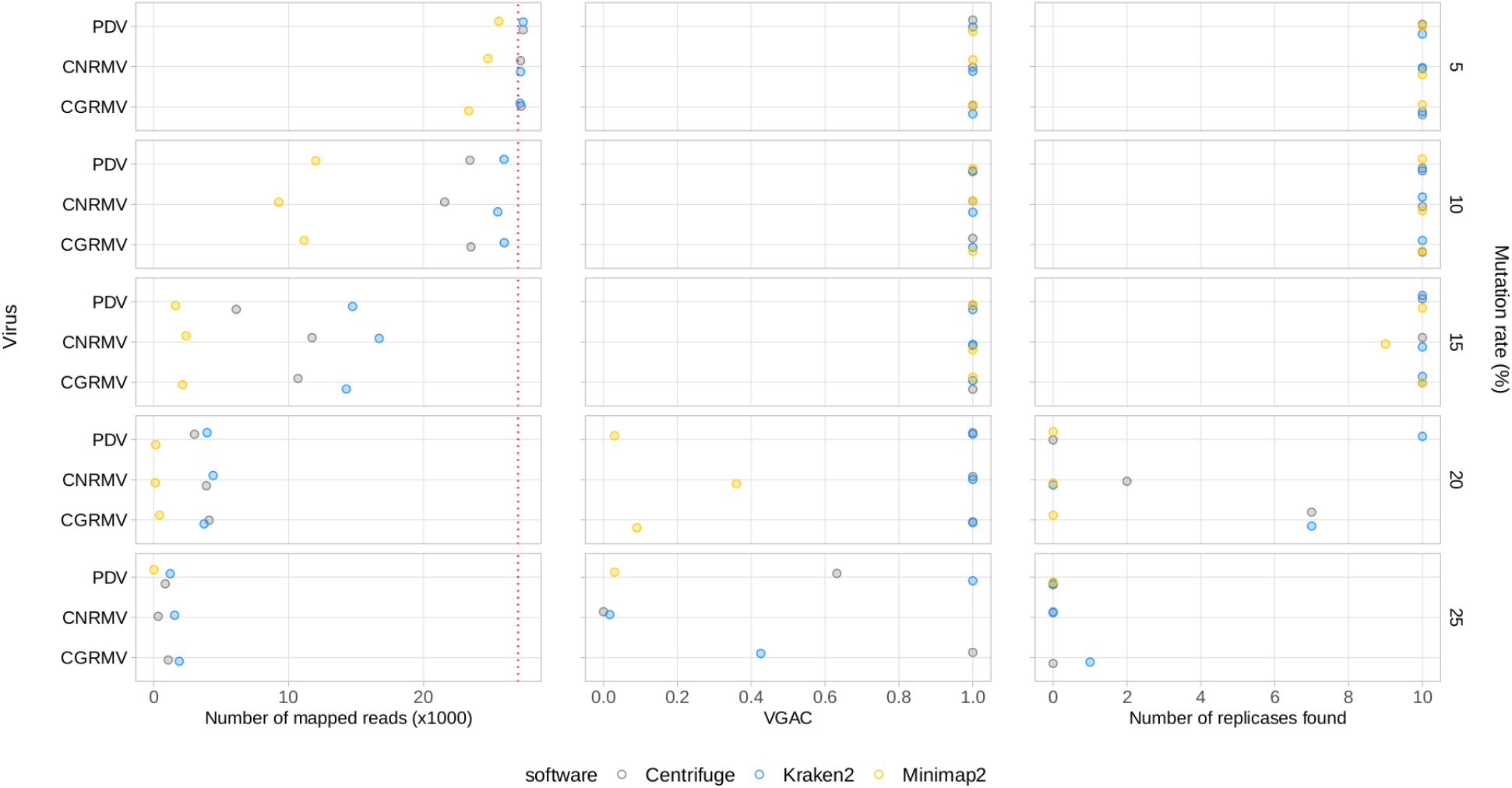
Simulated mutations in synthetic NGS data. Mutated virus genomes were simulated to generate synthetic NGS dataset to evaluate read assignment tolerance to viral mutated variants. Each viral genome of the Pavium panel-I was randomly mutated at the different rates indicated (5, 10, 15, 20 and 25%) at the far right of each chart. Albeit 20 million reads were generated for each mutation rate, a 10x subsampling of 10 million reads was performed, so dots represent a mean number of assigned reads. The distribution of viral reads in this case was homogeneous. The cases of viruses CGRMV, CNRMV, and PDV are shown, but a full version of this figure is depicted in Supplementary Figure S4. Dotted red line: expected number of assigned (mapped) reads according to the distribution of reads. Average values (dots) as well as standard deviations are listed in Supplementary Table S11.

### Assembly coverage, viral abundance and replicase identification

In order to explore beyond the presence of reads and to examine the biological relevance of the assemblies obtained at different sequencing depths and viral abundance, the VGAC was analyzed in terms of the presence of replicases in the assembled contigs. This was performed to understand whether the assembled portion of the virus could encode a relevant biological function to support the use of specific VGAC cutoffs for virus detection.

A series of simulated samples containing an increasing amount of viral reads with at most 3% of the total reads were generated (see Materials and Methods). In this case, only the viruses with the shortest and the longest genomes in the Pavium panel-I were surveyed (i.e. CVA and LChV-1, respectively). Simulated data (Cva and Lchv1 datasets) showed that all assessed software reached similar levels of VGAC values. At the lowest depth of sequencing (100K), virus abundance was critical since VGAC turned out to be relevant only from 5% (CVA and LChV-1) of the viral read composition (Figure 5). However, the VGAC value increased at lower viral abundance as the depth of sequencing increased. At the lowest viral abundance (0.05%), the maximum VGAC values obtained at 10M total reads were 0.97 in the case of CVA and 0.89 for LChV-1. In extreme scenarios (e.g. 0.05% of total viral reads) the depth of sequencing became critical. Thus, at 10M total reads, the number of viral reads was ~150 (enough to assemble the longest virus), but at 100K total reads, this number was ~1.5 reads, making the assembly of a viral genome not possible (thus VGAC ≈ 0).

**Figure 5.**
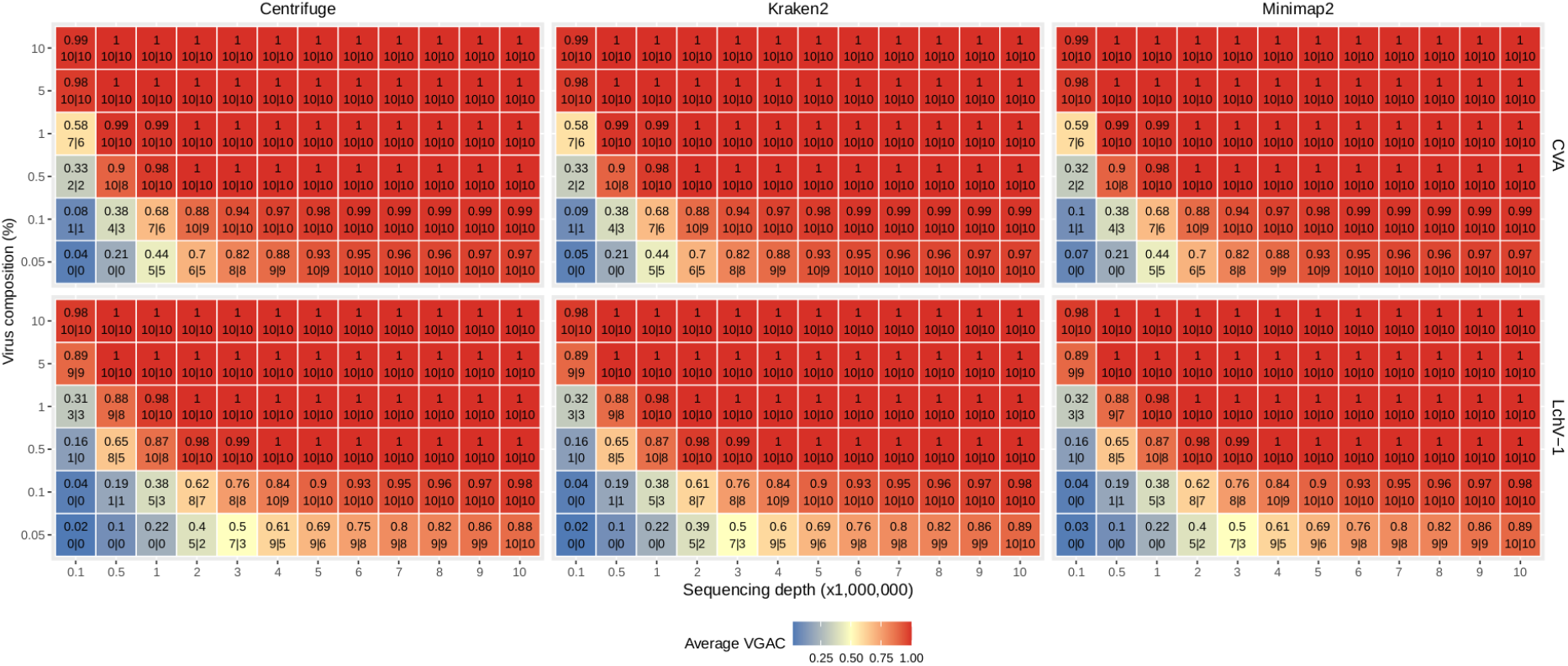
VGAC dependence on viral abundance, depth of sequencing and identification of replicases. In order to analyze the relationship of VGAC on viral abundance, depth of sequencing and the presence of replicases, synthetic NGS data were generated with different abundance of viral reads. The viruses with the shortest and the longest genome length of the Pavium panel-I were assayed only (CVA and LChV-1, respectively). The heatmap represents the VGAC obtained (indicated in each cell in the upper value) at both the respective sequencing depth and the abundance (%) of the viral reads (represented in the vertical axis as % viral composition of the corresponding 3% of viral reads of the sequencing depth); bottom numbers inside a cell indicates the number of replicases found in the respective assembly of the 10 subsampled sets at 80% alignment length / 80% similarity, and 90% alignment length /90% similarity (both numbers separated by a pipe symbol).

Further, a set of 17,708 viral proteins related to replication (e.g. replicase or polymerase) obtained from *RVDB-prot* (Bigot et al. 2020) was used to identify replicases in the assembled contigs. A minimum of 90% alignment length and 90% similarity between the assemblies and the set of replicases yielded a high correlation (R^2^ > 0.9) between the VGAC and the presence of replicases, which is expected as long as a full length virus can be assembled (Figure 5). When lowering the parameters to 80% alignment length and 80% similarity, there was a subtle increase in the recovery of replicases at lower depth of sequencing and lower viral abundance. At these parameters, correlation between the VGAC and the presence of replicases still remained high (R^2^ > 0.9).

Altogether, according to the simulations and independently of the software used, the identification of replicases began at a VGAC ≈ 0.3 for the lowest depth of sequencing case (100K total reads) or at VGAC ≈ 0.4 for the lowest viral abundance case (0.05%), which represents basically a minimum of 15 viral reads for CVA or 30 reads for LChV-1 required for being able to identify a replicase in some of the simulations (Figure 5). Coincidentally, the LChV-1 genome is 16,934 bp length and CVA is 7,383 bp length, which reflects the requirement of twice the number of reads for the identification.

Additionally, the presence of replicases was evaluated for the simulated mutation datasets, where the identification was expected to be affected according to the mutation rate since changes in the nucleotide sequence may alter the encoded protein. The identification of replicases was severely hampered over a 15% mutation rate (Figure 4 and Supplementary Figure S4), but a VGAC of 1 was still obtained, and even at 20% using *Centrifuge* and *Kraken2*. These results were the basis for establishing a 90% sequence identity threshold for clustering and to incorporate different isolates in a reference database used in the read assignment process (i.e. for building the Pavium panel-II).

### Performance on external published datasets

External datasets were used in order to challenge the methodology proposed in this study. Ten illumina datasets (V1-V10) comprising a panel of 18 viruses published by Tamisier et al. 2021 were used (Viromock datasets; Supplementary Table S4). In addition, three small RNA sequencing datasets (R1-R3) published by Barrero et al. 2017 and Jo et al. 2016, 2017 (SmallRNA datasets; Supplementary Table S4) were included in this analysis by generating a panel of 21 viruses.

According to the performance on Viromock datasets (Table 1 and Supplementary Table S7), the pipeline is able to achieve the detection of viral reads, to assemble a partial or full viral genome, and to detect replicases in 14 out of 15 cases under the criterion of detection with at least twoout of three software (93% in both schemas 80/80 and 90/90). The case V3-GRLaV2 did not meet the criterion since replicase detection was possible in the assembly from reads assigned only by one software (*Centrifuge*, in both schemas) due to lower VGAC values. In the same dataset the cases V3-GRSPaV and V3-GRVFV failed the detection with the reads assigned by *Minimap2*. In those three cases the VGAC values were less than 0.2, but replicases were detected above this value.

**Table 1.**
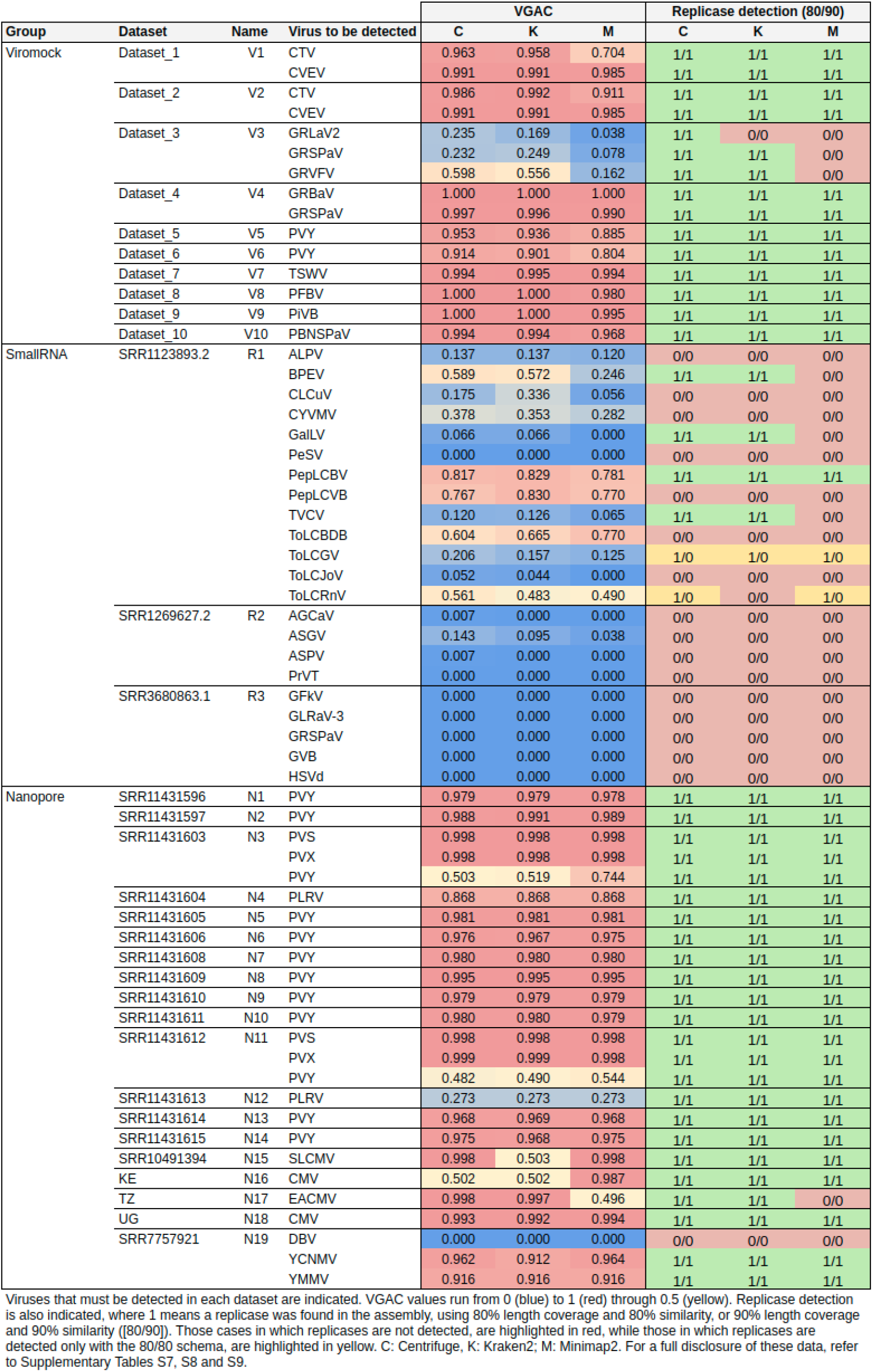
Summary of performance of pipeline on external datasets.

Regarding SmallRNA datasets (Table 1 and Supplementary Table S8), four cases could complete the pipeline (read-assignment, assembly, pseudo-annotation) with at least two out of three software (cases R1-PepLCBV, R1-BPEV, R1-GalLV, and R1-TVCV). The case of BPEV showed a low number of reads but enough to assemble a region to find a replicase (*Centrifuge* and *Kraken2*). The GalLV case can be considered artifactual since the actual replicase is encoded in the first 6,000 bp of the 5’-region of the reference genome (one segment of 8,659 bp length), but a shorter replicase was found at the end of the 3’-region (hence it was detected at very low VGAC values). In the case of TVCV, low VGAC values were obtained, however a contig could be assembled around position 4,300 bp of the reference genome (which is 7,767 bp length) containing the replicase. In the CLCuV case, there was a high number of reads, however they could only assemble a contig that maps in the central region of the reference genome, and the replicase in this virus is at the 3’-region (hence no replicases detected). A similar issue was observed in the case CYVMV, where no contig could be assembled for the 3’-region of the reference genome where the replicase is located. In the PepLCVB and ToLCBDB cases, despite the relatively high VGAC values, no replicases were found. This was expected since betasatellite viruses have been reported to depend entirely on other viruses for replication, movement, and transmission (Shafiq et al., 2020). In the ToLCJoV case, the pipeline could be completed only with the 80/80 schema, but it could be considered artifactual since the actual replicase of the virus is in the 3’-region, which could not be assembled (the assembled contigs map to the 5’-half of the reference genome, which contains a replicase encoded in the 3’-to-5’ direction). In the ToLCRnV case, the pipeline could detect replicases only at the 80/80 schema for *Centrifuge* and *Minimap2*. This is because the consensus contig for reads assigned by *Centrifuge* or *Minimap2* was longer than for reads assigned by *Kraken2*, thus contained a replicase.

No detection could be done in the datasets R2 or R3 despite reads being assigned with all software in the former dataset, and only with *Centrifuge* in the latter dataset. All these cases were manually inspected verifying that either assembly was not possible (e.g. due to length of reads) or, due to reads being localized in a region of the reference genome that did not contain at least 80% of a replicase (e.g. the case R2-ASGV for *Minimap2* depicted in Supplementary Figure S5).

Additionally, Nanopore sequencing datasets composed of 19 samples (N1-N19) published by Boykin et al. 2018, Della-Bartola et al. 2020, Filloux et al. 2018, and Leiva et al. 2020 (Nanopore datasets; Supplementary Table S4) were also subjected to the pipeline, including 10 target viruses for detection. According to the performance on these datasets (Table 1 and Supplementary Table S9), the pipeline was able to assign viral reads, to assemble a partial or full viral genome, and to identify replicases in 24 out of 25 cases (96%) using the criterion of two out of three software and both schemas (80/80 and 90/90), and 1 case with no read assignment (case N19-DBV). All the cases were also manually inspected, finding that some of the replicases were detected in assemblies with lower VGAC values. For example, in the cases N3-PVY (VGAC ≈ 0.5-0.7), N11-PVY (VGAC ≈ 0.5), and N12-PLRV (VGAC ≈ 0.3) the 3’-regions of the viruses could be assembled, which are the regions that encode their respective replicases; in the case N15-SLCMV (a 2-segments virus), reads assigned by *Kraken2* could assemble one of the segments of the virus (thus VGAC ≈ 0.5), which resulted to be the one encoding the replicase; in the case N16-CMV (a 2-segments virus), reads assigned by *Minimap2* could assemble both segments of the virus (thus VGAC > 0.9), and reads assigned by *Centrifuge* and *Kraken2* could assemble one segment (VGAC ≈ 0.5), nonetheless the three assemblies encoded the replicase (in this case, the number of reads was less than 60, although with ~3,200 bp length on average; Supplementary Table S4); finally, in the case N17-EACMV (a 2-segments virus), despite the number of reads assigned by *Minimap2*, one segment could be assembled (the one lacking the replicase; thus VGAC ≈ 0.5). The Dataset N19 was further investigated to confirm the lack of reads assigned to DBV. Although the dataset was reported to have 156 ONT reads (Filloux et al., 2018), it was not possible to assign them with the 3 software, nor additionally when using *Diamond* and *Blast* tools.

Overall performance on external datasets is presented in Table 2 (Supplementary Table S8). Since the datasets are reported to have only the viruses that can be detected (true positives), true negative and false positive cases were estimated from the rest of viruses of the respective panel which account for misassigned reads by at least one software (see Material and Methods). The three software performed with high sensitivity at the read assignment level (average > 0.9), but specificity was disparate amongst them with *Minimap2* reaching the highest degree (average = 0.82 in comparison with 0.26-0.36 reached by the other two software). Similar patterns, where *Minimap2* outperformed *Centrifuge* and *Kraken2*, were obtained for precision (0.81 vs 0.55-0.56) and accuracy (0.85 vs 0.61-0.60). This was in agreement with the FDR values, where the pattern was the opposite, wherein *Minimap2* showed the lowest rate (0.19 vs 0.45-0.44). Regarding the metrics for VGAC, sensitivity and accuracy values tended to be lower as long as cutoffs became more stringent, meanwhile specificity, precision, and FDR tended to improve reaching the maximal (1) and the minimal values (0), respectively. In relation to the detection of replicases, both schemas (80/80 and 90/90) appeared to have identical measure values in the Viromock and Nanopore datasets, while the use of either schemas appeared to have more impact in the SmallRNA datasets (for instance, the average sensitivity at the 80/80 schema was 0.21, while in the 90/90 schema, 0.14). Altogether, these measures provide additional rationale for the high performance of the Viromock (Illumina datasets) and Nanopore datasets (Table 2), in which the pipeline could completely identify more than 93% of the tested cases.

**Table 2.** Performance on external datasets.

| Group | Measure | Read Assignment | | | | VGAC $\geq 0.1$ | | | | VGAC $\geq 0.3$ | | | | 80/80 | | | | Distance | |
| --- | --- | --- | --- | --- | --- | --- | --- | --- | --- | --- | --- | --- | --- | --- | --- | --- | --- | --- | --- |
|  |  | C | K | M | Ave | C | K | M | Ave | C | K | M | Ave | C | K | M | Ave | VGAC0.1 | VGAC0.3 |
| Viromock | Sensitivity | 1.00 | 1.00 | 1.00 | **1.00** | 1.00 | 1.00 | 0.87 | **0.96** | 0.87 | 0.87 | 0.80 | **0.84** | 1.00 | 0.93 | 0.80 | **0.91** | 0.0943 | 0.1491 |
|  | Specificity | 0.24 | 0.24 | 0.86 | **0.44** | 1.00 | 1.00 | 1.00 | **1.00** | 1.00 | 1.00 | 1.00 | **1.00** | 1.00 | 1.00 | 1.00 | **1.00** | 0.0000 | 0.0000 |
|  | Precision | 0.28 | 0.28 | 0.68 | **0.41** | 1.00 | 1.00 | 1.00 | **1.00** | 1.00 | 1.00 | 1.00 | **1.00** | 1.00 | 1.00 | 1.00 | **1.00** | 0.0000 | 0.0000 |
|  | Accuracy | 0.41 | 0.41 | 0.89 | **0.57** | 1.00 | 1.00 | 0.97 | **0.99** | 0.97 | 0.97 | 0.95 | **0.96** | 1.00 | 0.98 | 0.95 | **0.98** | 0.0214 | 0.0339 |
|  | FDR | 0.72 | 0.72 | 0.32 | **0.59** | 0.00 | 0.00 | 0.00 | **0.00** | 0.00 | 0.00 | 0.00 | **0.00** | 0.00 | 0.00 | 0.00 | **0.00** | 0.0000 | 0.0000 |
| SmallRNA | Sensitivity | 1.00 | 0.77 | 0.77 | **0.85** | 0.50 | 0.45 | 0.36 | **0.44** | 0.27 | 0.32 | 0.18 | **0.26** | 0.27 | 0.23 | 0.14 | **0.21** | 0.3936 | 0.1016 |
|  | Specificity | 0.33 | 0.50 | 0.92 | **0.58** | 1.00 | 1.00 | 1.00 | **1.00** | 1.00 | 1.00 | 1.00 | **1.00** | 1.00 | 1.00 | 1.00 | **1.00** | 0.0000 | 0.0000 |
|  | Precision | 0.73 | 0.74 | 0.94 | **0.81** | 1.00 | 1.00 | 1.00 | **1.00** | 1.00 | 1.00 | 1.00 | **1.00** | 1.00 | 1.00 | 1.00 | **1.00** | 0.0000 | 0.0000 |
|  | Accuracy | 0.76 | 0.68 | 0.82 | **0.75** | 0.68 | 0.65 | 0.59 | **0.64** | 0.53 | 0.56 | 0.47 | **0.52** | 0.53 | 0.50 | 0.44 | **0.49** | 0.2547 | 0.0658 |
|  | FDR | 0.27 | 0.26 | 0.06 | **0.19** | 0.00 | 0.00 | 0.00 | **0.00** | 0.00 | 0.00 | 0.00 | **0.00** | 0.00 | 0.00 | 0.00 | **0.00** | 0.0000 | 0.0000 |
| Nanopore | Sensitivity | 0.96 | 0.96 | 0.96 | **0.96** | 0.96 | 0.96 | 0.96 | **0.96** | 0.92 | 0.92 | 0.92 | **0.92** | 0.96 | 0.96 | 0.92 | **0.95** | 0.0385 | 0.0544 |
|  | Specificity | 0.22 | 0.33 | 0.67 | **0.41** | 1.00 | 1.00 | 0.94 | **0.98** | 1.00 | 1.00 | 0.94 | **0.98** | 1.00 | 1.00 | 0.94 | **0.98** | 0.0000 | 0.0000 |
|  | Precision | 0.64 | 0.68 | 0.81 | **0.71** | 1.00 | 1.00 | 0.96 | **0.99** | 1.00 | 1.00 | 0.96 | **0.99** | 1.00 | 1.00 | 0.96 | **0.99** | 0.0015 | 0.0000 |
|  | Accuracy | 0.66 | 0.70 | 0.84 | **0.73** | 0.98 | 0.98 | 0.95 | **0.97** | 0.95 | 0.95 | 0.93 | **0.95** | 0.98 | 0.98 | 0.93 | **0.96** | 0.0227 | 0.0321 |
|  | FDR | 0.36 | 0.32 | 0.19 | **0.29** | 0.00 | 0.00 | 0.04 | **0.01** | 0.00 | 0.00 | 0.04 | **0.01** | 0.00 | 0.00 | 0.04 | **0.01** | 0.0015 | 0.0000 |
| Average All | Sensitivity | **0.99** | **0.91** | **0.91** | **0.94** | **0.82** | **0.81** | **0.73** | **0.79** | **0.69** | **0.70** | **0.63** | **0.68** | **0.74** | **0.71** | **0.62** | **0.69** | **0.1662** | **0.0594** |
|  | Specificity | **0.26** | **0.36** | **0.82** | **0.48** | **1.00** | **1.00** | **0.98** | **0.99** | **1.00** | **1.00** | **0.98** | **0.99** | **1.00** | **1.00** | **0.98** | **0.99** | **0.0000** | **0.0000** |
|  | Precision | **0.55** | **0.56** | **0.81** | **0.64** | **1.00** | **1.00** | **0.99** | **1.00** | **1.00** | **1.00** | **0.99** | **1.00** | **1.00** | **1.00** | **0.99** | **1.00** | **0.0005** | **0.0000** |
|  | Accuracy | **0.61** | **0.60** | **0.85** | **0.69** | **0.88** | **0.87** | **0.84** | **0.87** | **0.82** | **0.83** | **0.79** | **0.81** | **0.84** | **0.82** | **0.78** | **0.81** | **0.0955** | **0.0214** |
|  | FDR | **0.45** | **0.44** | **0.19** | **0.36** | **0.00** | **0.00** | **0.01** | **0.00** | **0.00** | **0.00** | **0.01** | **0.00** | **0.00** | **0.00** | **0.01** | **0.00** | **0.0005** | **0.0000** |
| Average Viromock + Nanopore | Sensitivity | **0.98** | **0.98** | **0.98** | **0.98** | **0.98** | **0.98** | **0.91** | **0.96** | **0.89** | **0.89** | **0.86** | **0.88** | **0.98** | **0.95** | **0.86** | **0.93** | **0.0622** | **0.1007** |
|  | Specificity | **0.23** | **0.28** | **0.76** | **0.43** | **1.00** | **1.00** | **0.97** | **0.99** | **1.00** | **1.00** | **0.97** | **0.99** | **1.00** | **1.00** | **0.97** | **0.99** | **0.0000** | **0.0000** |
|  | Precision | **0.46** | **0.48** | **0.74** | **0.56** | **1.00** | **1.00** | **0.98** | **0.99** | **1.00** | **1.00** | **0.98** | **0.99** | **1.00** | **1.00** | **0.98** | **0.99** | **0.0008** | **0.0000** |
|  | Accuracy | **0.53** | **0.56** | **0.87** | **0.65** | **0.99** | **0.99** | **0.96** | **0.98** | **0.96** | **0.96** | **0.94** | **0.96** | **0.99** | **0.98** | **0.94** | **0.97** | **0.0204** | **0.0326** |
|  | FDR | **0.54** | **0.52** | **0.26** | **0.44** | **0.00** | **0.00** | **0.02** | **0.01** | **0.00** | **0.00** | **0.02** | **0.01** | **0.00** | **0.00** | **0.02** | **0.01** | **0.0008** | **0.0000** |

Refer to Materials and Methods for more details on these measures.
Distance between measures of VGAC and 80/80 schema.
For a full disclosure of these data, refer to Supplementary Table S10.
C: Centrifuge; K: Kraken2; M: Minimap2; Ave: Average value

From these observations, the 80/80 schema appeared to be more suitable for the detection of replicases since it allows obtaining higher performance while still being composed of stringent thresholds (80% similarity and 80% alignment length). In that sense, a comparison of the measures obtained under these thresholds with the measures for VGAC at the different cutoffs, was carried out so as to find the cutoff at which similar performance measures are obtained with such a schema, accounting for the minimum VGAC in which replicases could be identified. To identify this VGAC cutoff, the euclidean distance between the measures (sensitivity, specificity, accuracy, precision, and FDR) was calculated (see Material and Methods). A minimal of the distances was found at VGAC ≥ 0.2 for Viromock datasets, VGAC ≥ 0.4 for SmallRNA datasets, VGAC ≥ 0.1 for Nanopore datasets, and VGAC ≥ 0.3 taking into account all the datasets (Figure 6).

**Figure 6.**
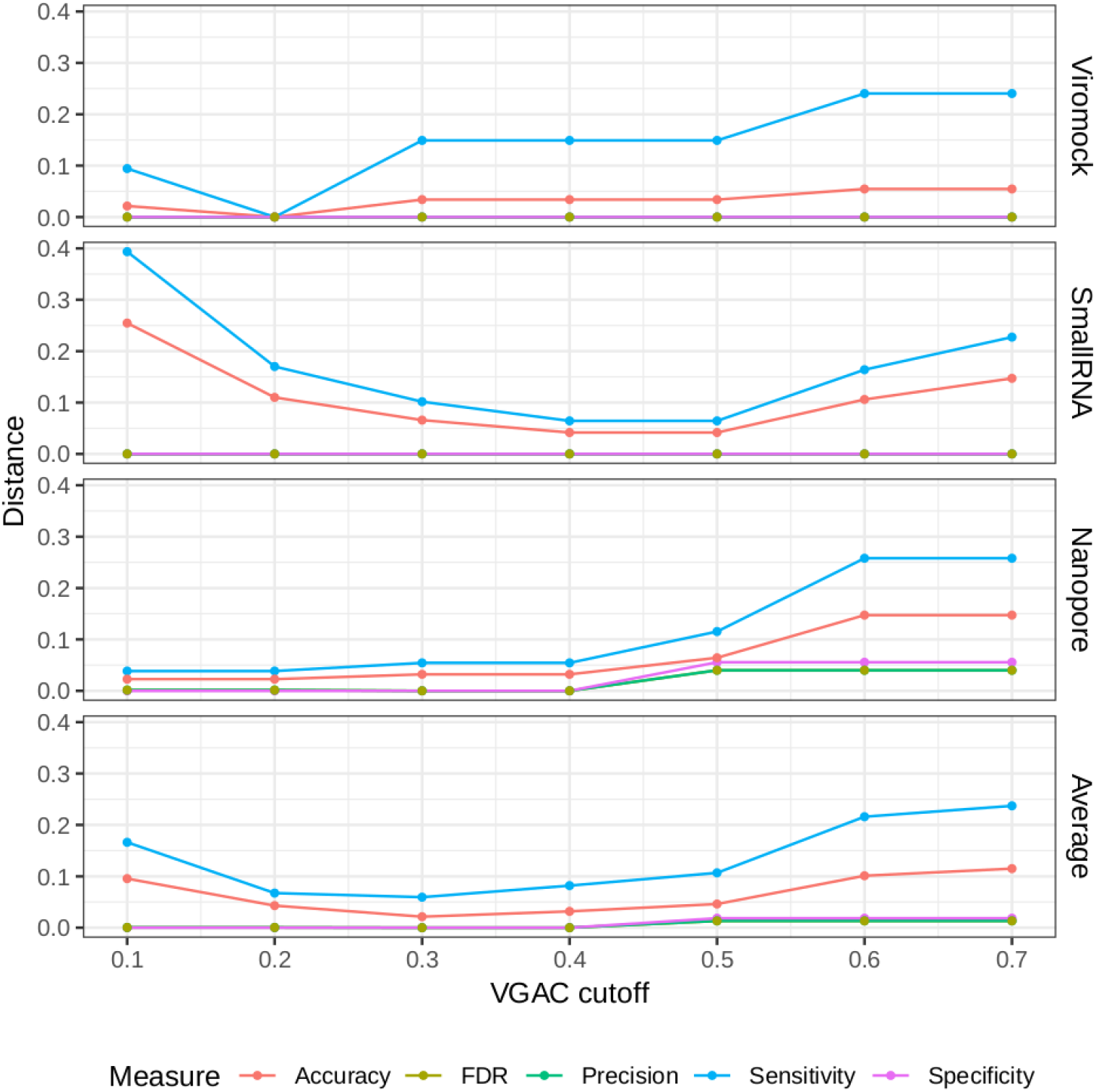
Distances between measures from external datasets. Comparison of the measures (accuracy, FDR, precision, sensitivity and specificity) obtained under different cutoffs of VGAC with the measures obtained for the identification of replicases under the 80/80 schema. Comparison was carried out in term of the distance between the measures according to Materials and Methods. For a detailed description refer to Supplementary Table S10.

### Comparison of Viroscope pipeline and RT-PCR analysis

All sweet cherry samples used in this study were tested by external and internal RT-PCR methods (see Materials and Methods) to confirm the presence or absence of the 11 viruses previously evaluated by Illlumina sequencing (Table 3 and Supplementary Figure S6). Additionally, seasonal effects were also assessed. Samples for the same plant specimens were collected at spring (SD-L1, SD-L2, SD-S1, and SD-S2, from the shoot development stage) and at the end of summer (SS-L1, SS-L2, SS-S1, and SS-S2, from the senescence stage). A total of 88 analyses by RT-PCR (external and internal) account for the detection of the 11 viruses. The summary of the results including the internal PCR, the external PCR, and the virus detection through Viroscope is shown in Table 3. The diagnosis performed by Viroscope used the following criteria based on the results aforementioned: a VGAC ≥ 0.3 for a positive virus detection; a VGAC between 0.1 and 0.3 for a positive virus detection only when a replicase can be identified; a negative detection for samples with VGAC <0.1; finally a requisite of agreement between two of the three software used. In this case, the Pavium panel-II was used as a reference database (see Materials and Methods). The detailed results of the diagnosis conducted by the pipeline are depicted in Figure 7 (shoot development stage, SD) and Figure 8 (senescence stage, SS).

**Table 3.**
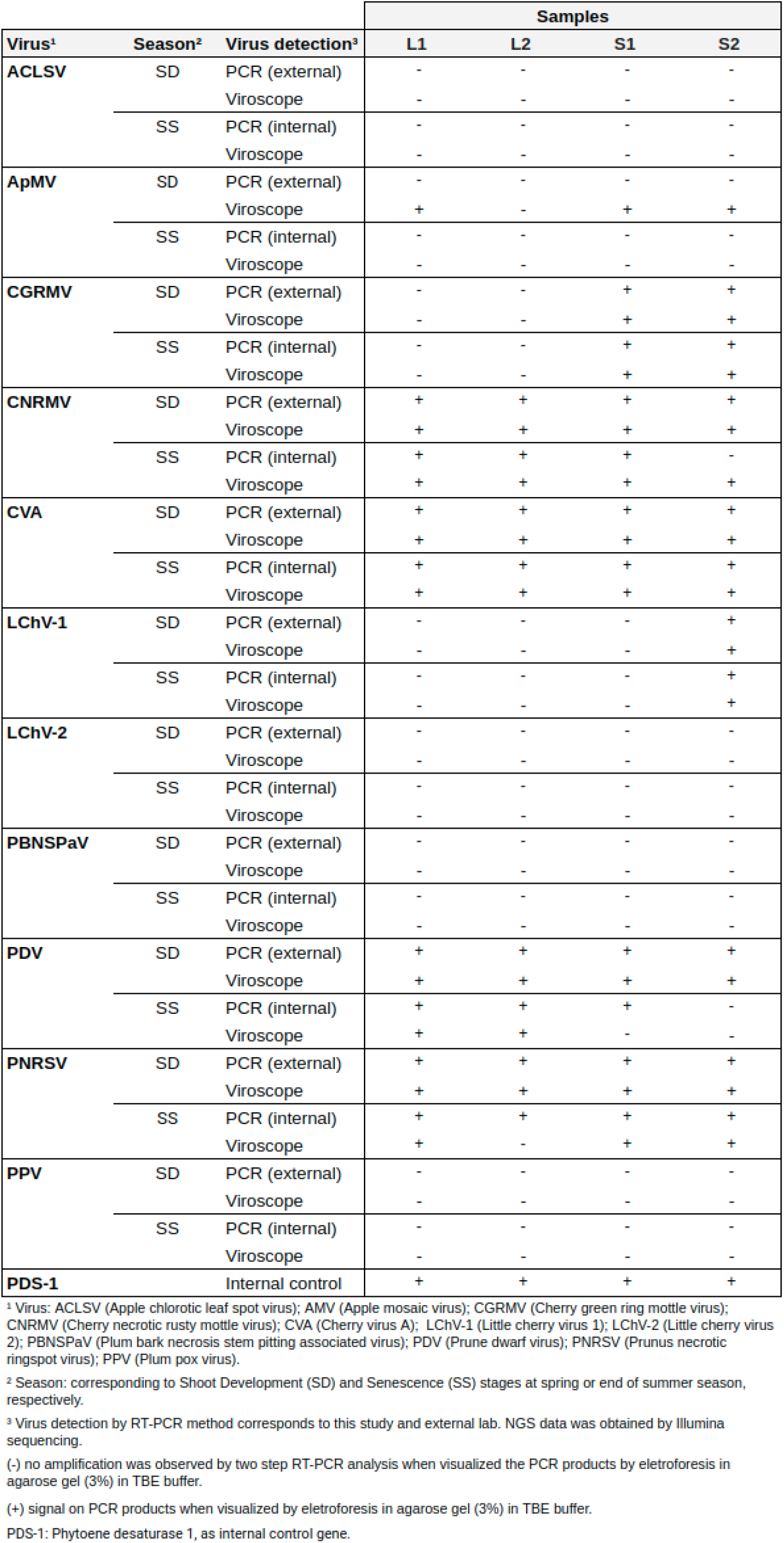
Pipeline validation by RT-PCR.

**Figure 7.**
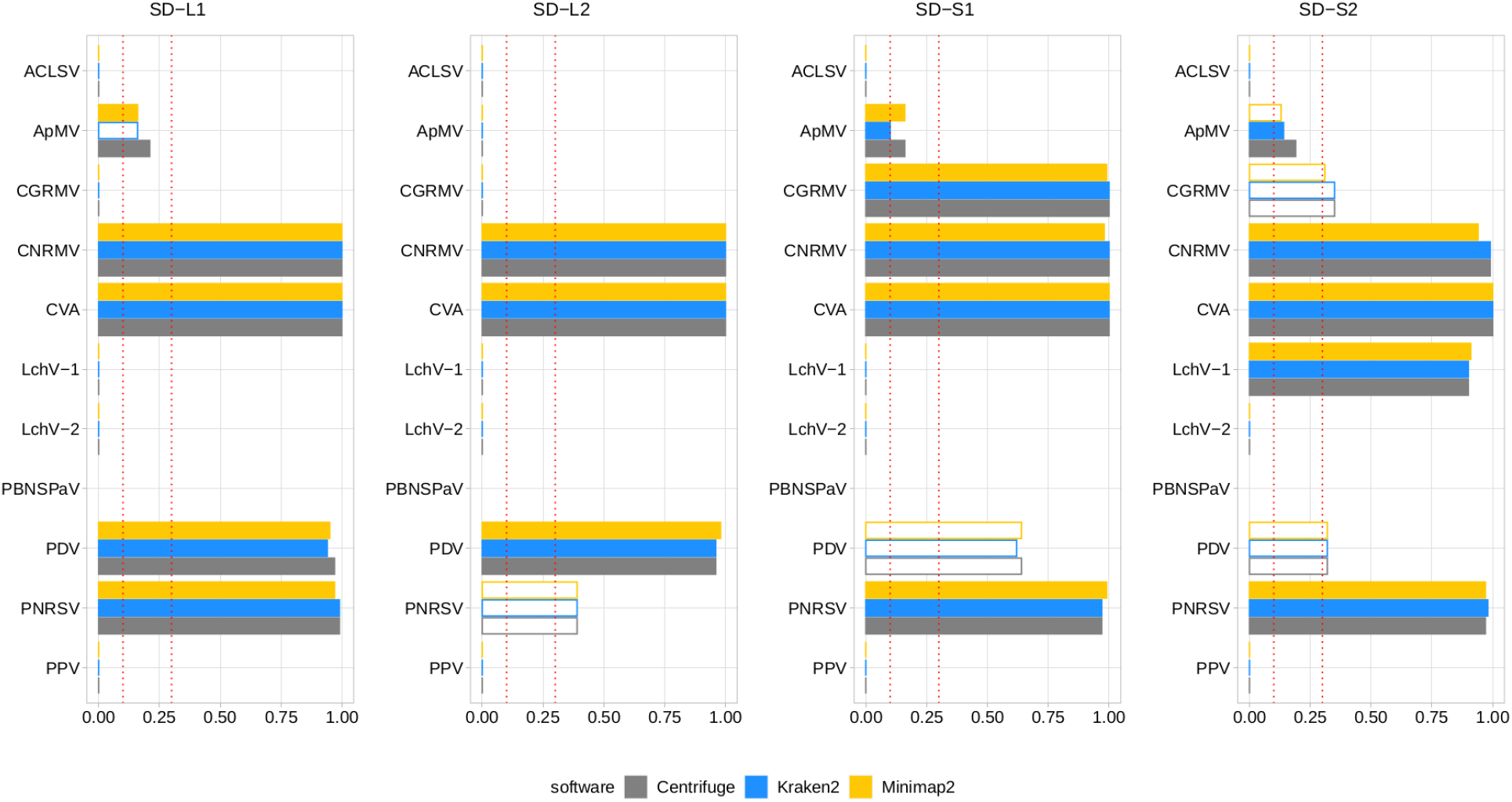
Diagnosis on field samples from the shoot develpoment stage using Pavium panel-II. NGS data from field samples (SD-L1, SD-L2, SD-S1, and SD-S2) were submitted to the pipeline using the Pavium panel-II. X-axis: VGAC scale (0 to 1); ordinate: abbreviated name of viruses; dotted lines: VGAC cutoffs at 0.1 and 0.3; filled bars: replicase identified. Supplementary Figure S7 depicts these data together with read assignment levels.

**Figure 8.**
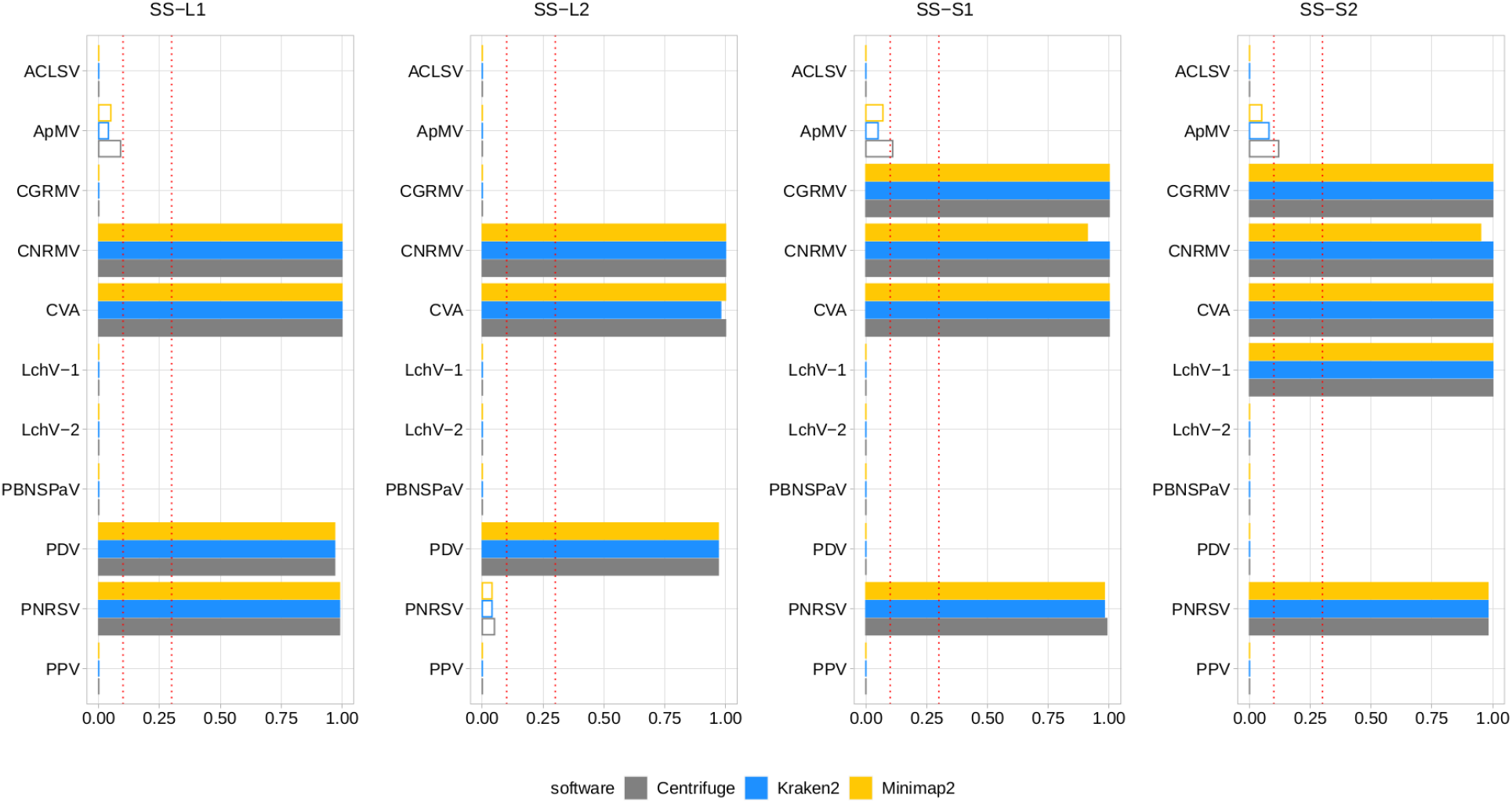
Diagnosis on field samples from the senescence stage using Pavium panel-II. NGS data from field samples (SS-L1, SS-L2, SS-S1, and SS-S2) were submitted to the pipeline using the Pavium panel-II. X-axis: VGAC scale (0 to 1); ordinate: abbreviated name of viruses; dotted lines: VGAC cutoffs at 0.1 and 0.3; filled bars: replicase identified. Supplementary Figure S8 depicts these data together with read assignment levels.

High consistency between NGS-based detection and RT-PCR analyses was observed, where 82 out of 88 (93.2%) matches were obtained. The inconsistencies observed between the RT-PCR analyses were detected only for sample S2, which was found positive for the CNRMV in the SD stage by the external PCR and negative in SS for the internal PCR, which was also the case for PDV. Regarding the Viroscope pipeline, CNRMV resulted in a positive case for both samples, that is, independently of the season conditions. On the other hand, the diagnosis by the pipeline agreed with each RT-PCR test in the case of PDV, accounting for the seasonality of the sampling. Contrasting the RT-PCR results obtained in the SD stage and the SS stage, some discrepancies were found (2 out of 44 cases; 4.5%): CNRMV in sample SS-S2 and PDV in sample SS-S2. These results suggest that sampling season affects the sensitivity of diagnosis, particularly having increased sensitivity in the shoot development stage.

Noticeably, the diagnosis by Viroscope appeared to be more sensible than RT-PCR methods (internal and external), since samples SD-L1, SD-S1 and SD-S2 resulted to be positive for ApMV (Figure 7), while none of the laboratories were able to detect it (Table 3). This was more evident considering the number of reads assigned by at least two of the three software, which ranged from 10K to 20K reads in the SD-L1 and SD-S2 samples, and 1.5K to 2.5K reads in the SD-S1 sample. Replicases were also identified by two of the three algorithms with VGAC between 0.1 and 0.3. Furthermore, the diagnosis for CNRMV in sample SS-S2 was positive according to Viroscope (VGAC = 1 and with identification of replicases), whereas for PDV (sample SS-S1) and PNRSV (sample SS-L2) were negative (VGAC = 0 in both cases). Although the number of reads for PNRSV (sample SS-L2) reached over 4K reads assigned by two out of the three algorithms, they were not able to assemble a contig with a VGAC over 0.1, nor that contained a replicase (Supplementary Figura S5). The specific cases of CNRMV, PDV and PNRSV point towards a difference of sensitivity or issues of amplification likely due mismatches in the primer binding region (false negative), or cross-contamination during sample as well as reagents manipulation (false positive). In fact, for PDV or PNRSV, the inspection of sequences showed that the annealing regions contained at least one mismatch to the primers used.

Comparing only the seasonal differences in diagnosis obtained through Viroscope, 6 cases were detected in SD and not in SS: ApMV in L1, L2 and S1; PDV in S1 and S2, and PNRSV in L2. For all these cases VGAC was below the minimum threshold and assigned reads dropped dramatically between both seasons. Furthermore, in the cases of SD-S1 and SD-S2 for PDV and SD-L2 for PNRSV were diagnosed as positive due to VGAC > 0.3, but no replicases were identified. These results again show that the shoot development stage enables increased sensitivity for virus detection using NGS-based diagnosis.

Lastly, it is important to highlight the use of different viral panels in the diagnosis process. According to the results described so far, VGAC values were not enough to diagnose LChV-1 as positive in sample SD-S2 when using the Pavium panel-I (Supplementary Figure S2), whereas appearing positive when using Pavium panel-II (Figure 7). The analysis of sequences revealed that the reference genome of LChV-1 in the first panel shares a 76% of sequence identity with the respective reference genome selected in the second panel, which explains the differences in the diagnosis.

### Web Application

The pipeline described in this study was implemented as a web application, *Viroscope*, to enable diagnosis of viruses on-demand from NGS data (accessible at https://www.viroscope.io). Users submit NGS data and then create an analysis instance for the pipeline to work and handle such data. Next, a predefined viral panel is selected to perform virus detection. Once finished, the pipeline outputs graphical results composed of a positivity report and taxonomical profile accounting for the abundance of each virus of the panel in the sample (Figure 9). The identification of the viruses is based on a read assignment by at least two of three algorithms (*Kraken2, Centrifuge* and *Minimap2*) and detection cutoff parameters are provided for the user to define, although suggestions are made according to the results obtained in this study. Three possible outcomes can be obtained from the pipeline: i) a categorically positive diagnosis which is defined when Viroscope was able to detect either the presence of a replicase or a VGAC over the upper cutoff defined by the user, ii) a categorically negative result which implies that read assignment was below the lower threshold of VGAC required for virus detection and with absence of replicase; and iii) a positive* (positive depicting an asterisk) result which establishes that read assignment resulted in a VGAC sufficient for virus detection but lower than what is required to attribute biological functionality. We surmise that lack of evidence for viral replication in these cases requires confirmation by other traditional methods to ensure diagnostic certainty.

**Figure 9.**
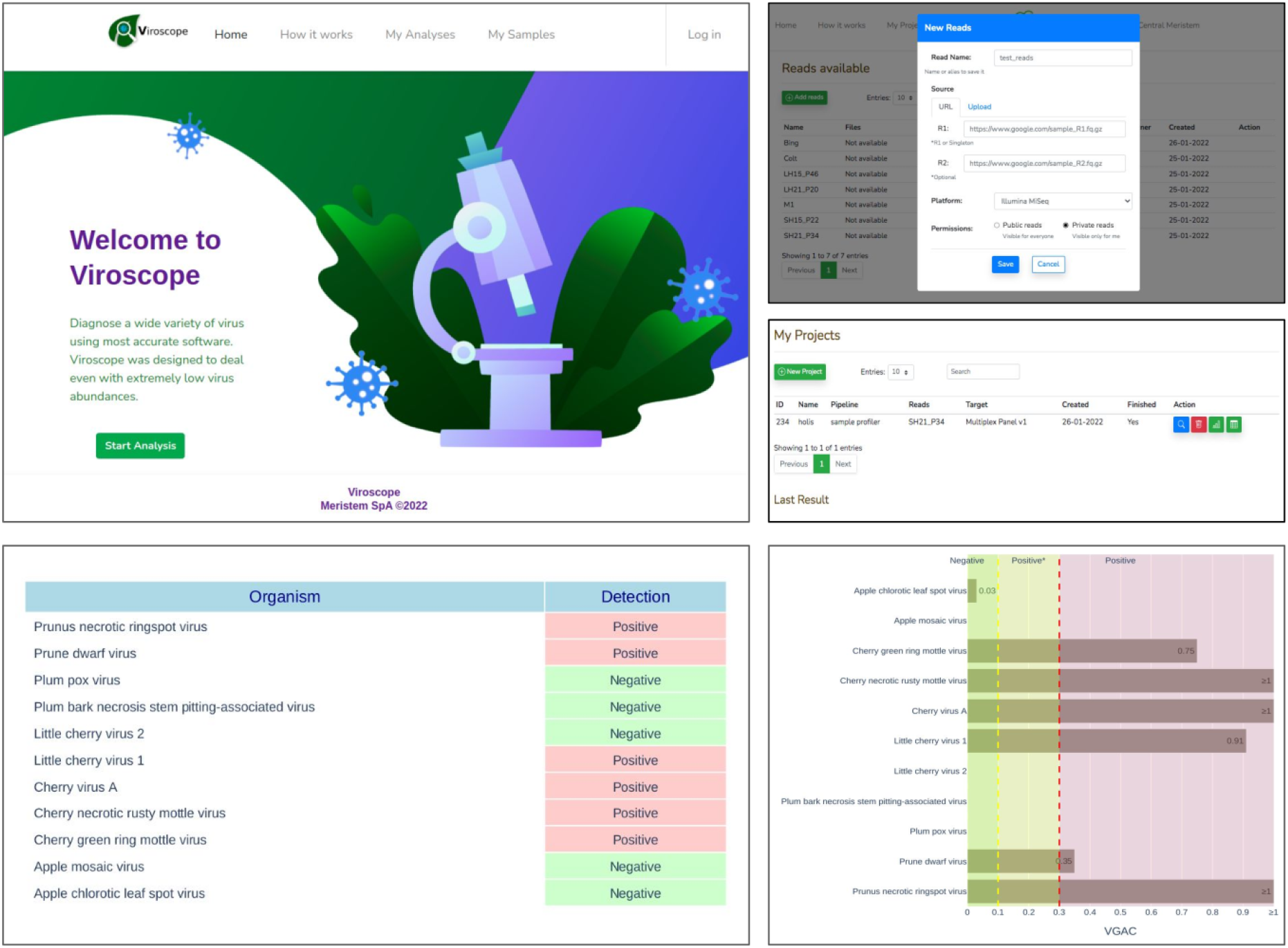
Viroscope.io. The Viroscope web application enables virus plant diagnostics using NGS data. It provides a user-interface to perform the Viroscope pipeline, and yields the VGAC metric as well as the presence of replicases to interpret the data and perform diagnosis. The process involves uploading NGS reads into the read library, selecting an analysis against a specific viral panel using user-inputted thresholds. This generates the analysis which is processed in real-time. The analysis then displays the results in plots that provide the metrics and suggests the diagnosis according to the selected thresholds, which is then confirmed by the user.

## DISCUSSION

In the last two decades, NGS methods have become a reliable tool for plant virus diagnostics including managing disease risk, emergence, and the adoption of novel phytosanitary rules (Adams et al., 2018; Gauthier et al., 2022). Due to its untargeted nature, NGS is capable of detecting multiple viruses (known as well as emergent ones) in infected material also when viruses are present in very low concentrations (Hanafi et al., 2022). In addition, it has proved to be a major advance for crops, imported plants and germplasm in which disease symptoms are absent, unspecific or only triggered by multiple viruses (Massart et al., 2017; Mehetre et al., 2021). In face of a changing farming paradigm, more efforts on NGS data analysis and its accurate interpretation must be established.

In this work, we developed Viroscope, a NGS data pipeline for virus diagnosis in plants that uses the VGAC and replicase identification to improve virus detection. One of the main challenges of virus diagnosis from NGS data is to define the virus viral presence when faced with a limiting amount of viral reads. Further, detection of a pathogen does not necessarily imply ongoing infection (Kiselev et al., 2020). We surmise that detection of viral sequences along with their capability to encode fundamental biological functions, such as replication, can improve certainty of detection by evaluating a functional aspect of virus biology.

The tests to evaluate metrics showed that VGAC is a more robust measure than read assignment for virus detection. Read assignment can be prone to generate false positives in circumstances where the host has sequences derived from viral origin, such as endogenous viral elements (Massart et al., 2019). Furthermore, the number of reads are directly influenced by sequencing depth and library preparation bias, and direct comparisons between datasets require normalization. In contrast, the use of VGAC is a better unifying measure despite the sequencing technology used, as depth of sequencing can differ in several orders of magnitude between platforms and laboratories, whilst VGAC is the result of an assembly, accounting for read length as well. VGAC rather than sequencing coverage demonstrated to be a better metric, as the latter can exhibit artificially high numbers when reads cover only a small portion of the genome or when contigs correspond to different viral variants, which can also exacerbate detection.

In turn, replicase identification from the assembled contigs enabled investigating the biological implication of different VGAC thresholds. Defining cutoffs for virus detection is extremely challenging and involves selecting a value that maximizes both sensitivity and specificity. We observed that replicases could be consistently identified only over a VGAC of 0.3, allowing us to define this threshold empirically as a level that enables a biological interpretation of the metric. Additionally, using replicase presence (or any other essential viral function) is already an excellent criteria for virus diagnosis in cases of even lower VGAC values.

Validation against external datasets shows that Viroscope has high diagnostic sensitivity and specificity for total RNA sequencing data. In contrast, small RNA data showed poor performance for virus detection with VGAC. Additionally, Nanopore sequencing data was used to evaluate if our pipeline is applicable to long-reads, where a 96% of agreement was obtained in comparison with the original publications (Boykin et al., 2018; Della-Bartola et al., 2020; Filloux et al., 2018; and Leiva et al., 2020). Long-read sequencing platforms have an advantage over short-read sequencing for VGAC as, regardless of the lower base calling quality, longer reads enable the complete reconstruction of a viral genome with fewer reads (e.g. CMV in sample N16). The overall performance is in agreement with the type of dataset analyzed, that is, the use of small RNA data appears to have a major impact on the sensitivity and the accuracy when VGAC or the replicase identification are concerned. This also reflects the fact that a greater VGAC cutoff is needed to obtain replicases, in contrast to Nanopore or the Illumina total RNA reads for which lower cutoffs were reported. Certainly, VGAC values are affected by the ability of the pipeline to assemble viruses (with the correct read assignment), hence the higher cutoffs in that type of data. In any event, the use of the VGAC allows to reach the maximum specificity and the minimum FDR even with VGAC ≥ 0.1, accounting for the elimination of false positives and false negatives from the minimum VGAC cutoff tested, being particularly evident in the performance for the Viromock and Nanopore datasets. Taking into account only total RNA sequencing data (Viromock and Nanopore datasets), a VGAC cutoff of ≥ 0.1 results in an average sensitivity of 96%, a specificity 99% and a FDR of 1%, which shows that Viroscope provides reliable detection.

Analysis of field samples by sequencing and RT-PCR showed that NGS-based virus diagnosis is more sensitive than RT-PCR and that seasonality of sampling influences viral abundance. Comparisons between RT-PCR panels showed that for both spring and summer samples, virus detection by RT-PCR had a 93.2% correspondence to NGS-based detection using Viroscope, where differences were due to false negative cases and false positive cases of detection by RT-PCR. Additionally, problems with PCR design bias were evident in one case in which primers contained mismatches to a viral variant. Further, the seasonality of sampling affected the capability to detect 6 viruses in summer through NGS. However, all viruses that could not be detected (ApMV, PDV, PNRSV) were Illarviruses, a genus named after exhibiting lability and described to be thermolabile (Hull, 2004). The cases of PDV and PNRSV are particularly interesting since no replicases were identified in the SD stage. For PNRSV, an approximate of 4K reads were assigned, but this did not produce an assembly with sufficient VGAC to be diagnosed as positive. Reads were mapped to a specific region in the 5’ end of one of the RNA segments of the viral genome. Possible explanations for this include endogenous viral element, an overcome infection, or host sequences being mapped to the virus. Nevertheless, this case and the ApMV cases emphasize the need to better understand viral physiology and the use of essential viral functions during detection for a certain diagnosis.

Viroscope uses a predefined panel of viral targets to perform diagnosis. This is due to the intended use of Viroscope as a viral diagnosis pipeline for phytosanitary detection rather than for discovery. We found that compiling a database of viral sequences using published reference sequences was insufficient to provide identification, as databases are generally biased toward submissions from certain geographical locations or particular pathogens. One such case occurred in the detection of LChV-1, as it could be identified by both PCR panels but not through Viroscope (using Pavium panel-I). The inclusion of other viral isolates to generate a clustered database (Pavium panel-II) enabled the successful identification through the pipeline. Despite this improvement in the diagnosis (if read assignment is concerned), more misassigned cases could be obtained as a side effect. The pipeline resulted to be sufficiently robust when VGAC and/or the identification of replicases are taken into account as misassigned reads were unable to assemble contigs. This finding is of paramount importance when considering performing the diagnosis in particular geographic contexts since viral isolates could be endemic or affect specific plant varieties, thus the panel construction should consider these distinctive features. Moreover, the viral variants can be dissimilar enough to have an impact on the diagnosis. Noticeably, the isolate included in the Pavium panel-II that allowed the identification of LChV-1 shares a 76% of sequence identity with the respective reference genome present in the Pavium panel-I.

NGS is now considered as the gold standard in molecular diagnostics of viral infections since it is a universal technique which is more precise at profiling pathogens (Rott et al., 2017; Massart et al., 2019; Kiselev et al., 2020; Soltani et al., 2021; Ruiz-García et al., 2021; Mehetre et al., 2021). As we have discussed in this work, plant NGS-based virus diagnostics is still facing many challenges related to both the analysis (standardization of metrics, harmonization of cutoff thresholds, biological interpretation), and at the regulatory-level implementation (detection of unregulated pathogens, ease-of-use and adoption). This work addresses some of the issues regarding the bioinformatics analysis and interpretation which we believe can aid in the ongoing discussion about how to implement these methods in real-world scenarios. Substantial research is still required to evaluate biological aspects of virus biology, particularly in cases of low abundance and low VGAC. Functional annotation may aid in increasing certainty of diagnosis in virus detection in these cases of low abundance, due to its direct implication in viral physiology and potential infectivity. In agreement with many scientific researchers in the field (Rott et al., 2017; Adams, 2018; Jones and Naidu, 2019; Gauthier et al., 2019; Mehetre et al., 2021; Villamor et al. 2021), we endorse that plant pathogen molecular diagnostics by NGS is becoming a more cost-effective and scalable solution with new accessible sequencing technologies already available to perform precise diagnosis. The enormous benefit of NGS applied to plant health is indispensable to ensure reliable diagnosis of known and unknown pathogens and will contribute to more sustainable agriculture and safer international plant trade practices.

## AVAILABILITY

Sequencing data are available at https://www.ncbi.nlm.nih.gov/sra/?term=PRJNA848006 Viroscope service is avaliable at https://www.viroscope.io

## SUPPLEMENTARY DATA

Supplementary Figures are available in PDF format and Supplementary Tables are avaliable in xlsx format in a zipped file.

## ACKNOWLEDGMENTS

We would like to thank Katerina Ferrat for coordinating sample submission and high-throughput sequencing. Alvaro Cuevas for field sampling of sweet cherry trees. Professor Dr. Juan Pablo Zoffoli, Cristian Yañez, and Dr. Victoria Cepeda for critically reading the manuscript and providing essential feedback.

