## Supplementary Figures for "Viroscope: plant viral diagnosis from NGS data using biologically-informed genome assembly coverage"

#### SUPPLEMENTARY FIGURE LEGENDS

**Figure S1. Read assignment with NGS data from field samples (full version).**
From sequencing data for field samples of cherry plants at the shoot development stage (SD-L1, SD-L2, SD-S1, and SD-S2), 10 subsets of randomly selected reads were built at different depths of sequencing. Three bioinformatic algorithms were tested, namely *Centrifuge*, *Kraken2*, and *Minimap2*. All the dotsbars represent the average of 10 measures. Read assignment at different depths of sequencing (note the different scales of the ordinates) using the Pavium panel-I (11 viruses). Average values (dotsbars) as well as standard deviations are listed in Supplementary Table S11.

**Figure S2. Viral genome assembly coverage with NGS data from field samples (full version).**
From sequencing data for field samples of cherry plants at the shoot development stage (SD-L1, SD-L2, SD-S1, and SD-S2), 10 subsets of randomly selected reads were built at different depths of sequencing. Three bioinformatic algorithms were tested, namely *Centrifuge*, *Kraken2*, and *Minimap2*. All the dotsbars represent the average of 10 measures. VGAC was calculated according to Materials and Methods and with the reads assigned by the different algorithms at the respective depth of sequencing. VGAC obtained from the assembly of assigned reads (ranges from 0 to 1). The cases for the 11 viruses form Pavium panel-I are presented. Average values (dotsbars) as well as standard deviations are listed in Supplementary Table S11.

**Figure S3. Read assignment and coverage in synthetic data (full version).**
Synthetic NGS data were generated to simulate actual samples. The Pavium panel-I comprising 11 viral genomes was used to build samples containing 3% of viral reads from a 20 million paired-end reads. Samples were built with both a homogeneous distribution of viral reads (*synhom*) and based on an average distribution of actual samples (*synab*) (Supplementary Table S2 and Supplementary Table S3). In all cases, 10 subsets of randomly selected reads were built at different depths of sequencing. Assigned reads are shown in the two first column charts and VGAC is shown in the last two column charts. Average values (dotsbars) as well as standard deviations are listed in Supplementary Table S11.

**Figure S4. Simulated mutations in synthetic NGS data (full version).**
Mutated virus genomes were simulated to generate synthetic NGS dataset to evaluate read assignment tolerance to viral mutated variants. Each viral genome of the Pavium panel-I was randomly mutated at the different rates indicated (5, 10, 15, 20, 25 and 30%) at the far right of each chart. Albeit 20 million reads were generated for each mutation rate, a 10x subsampling of 10 million reads was performed, so dotsbars represent a mean number of assigned reads. The distribution of viral reads in this case was homogeneous. Dotted red line: expected number of assigned (mapped) reads according to the distribution of reads. Average values (dotsbars) as well as standard deviations are listed in Supplementary Table S11.

**Figure S5. Examples of mapped reads onto viral genomes.**
**A**. Representation of ASGV genome (6,496 bp length). **B**. Assigned reads mapped onto ASGV (dataset R2), showing they were unable to assemble a contiguous contig to contain a replicase. **C**. Representation of segment 2 (2,046 bp length) of PNRSV genome. **D**. Assigned reads mapped onto segment 2 of PNRSV (sample SS-L2), where reads were concentrated almost only in the 5'-region. In this case, there were not reads mapping onto the other two segments (replicase is contained in segment 1).

**Figure S6. Viral pathogen detection in different infected plants of *Prunus sp.* according to the panel.**
A two-step singleplex RT-PCR analysis for the 11 viruses (lane 1–11) and the internal control (lane 12) was performed using specific primers. Leaf samples: (A) SS-L1; (B) SS-L2; (C) SS-S1; (D) SS-S2. The specific amplification products of the 11 viral pathogen (E) and the corresponding not template control (NTC) for each RT-PCR reaction (F) are shown. Lane 1: ACLSV; lane 2: AMV; lane 3: CGRMV; lane 4: CNRMV; lane 5: CVA; lane 6: LChV-1; lane 7: LChV-2; lane 8: PBNSPaV; lane 9: PDV; lane 10: PNRSV; lane 11: PPV and lane 12: PDS-1. MM: 100 bp molecular weight marker. The non-specific bands observed in sample S1 (C; lane 3) are not interfering with the expected PCR fragment for the virus CGRMV which is 181 bp long.

**Figure S7. Diagnosis on field samples from the shoot development stage using Pavium panel-II.**
NGS data from field samples (SD-L1, SD-L2, SD-S1, and SD-S2) were submitted to the pipeline using the Pavium panel-II. Left ordinate: number of viral reads (different scales); right ordinate: VGAC scale; circles: VGAC values for each software; filled circles: replicase identified; dotted lines: VGAC cutoffs at 0.1 and 0.3.

**Figure S8. Diagnosis on field samples from the senescence stage using Pavium panel-II.**
NGS data from field samples (SS-L1, SS-L2, SS-S1, and SS-S2) were submitted to the pipeline using the Pavium panel-II. Left ordinate: number of viral reads (different scales); right ordinate: VGAC scale; circles: VGAC values for each software; filled circles: replicase identified; dotted lines: VGAC cutoffs at 0.1 and 0.3.

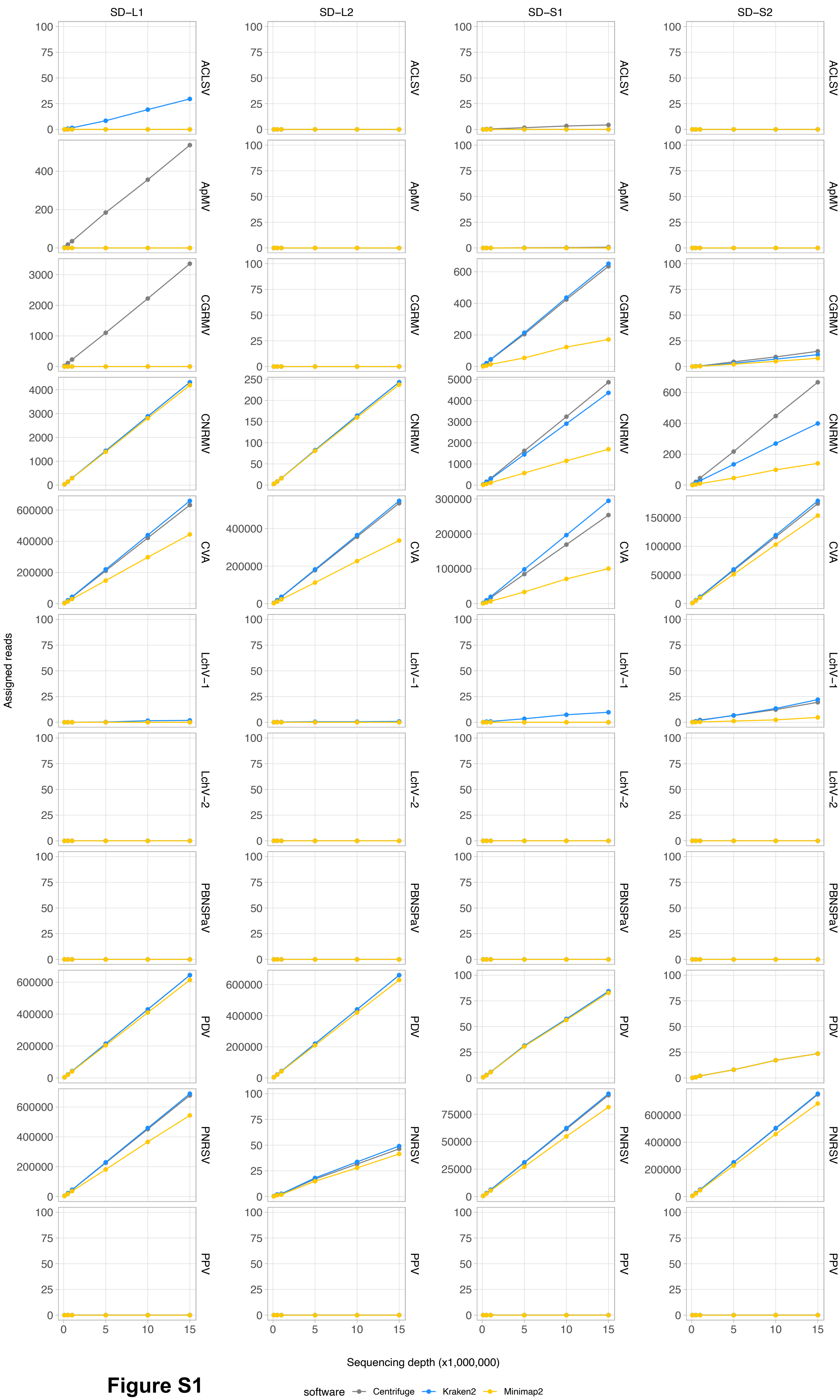

SD–L1
SD–L2
SD–S1
SD–S2
ACLSV
ApMV
CGRMV
CNRMV
CVA
LchV–1
LchV–2
PBNSPaV
PDV
PNRSV
PPV
Assigned reads
Sequencing depth (x1,000,000)
software
Centrifuge
Kraken2
Minimap2


**Figure S1**

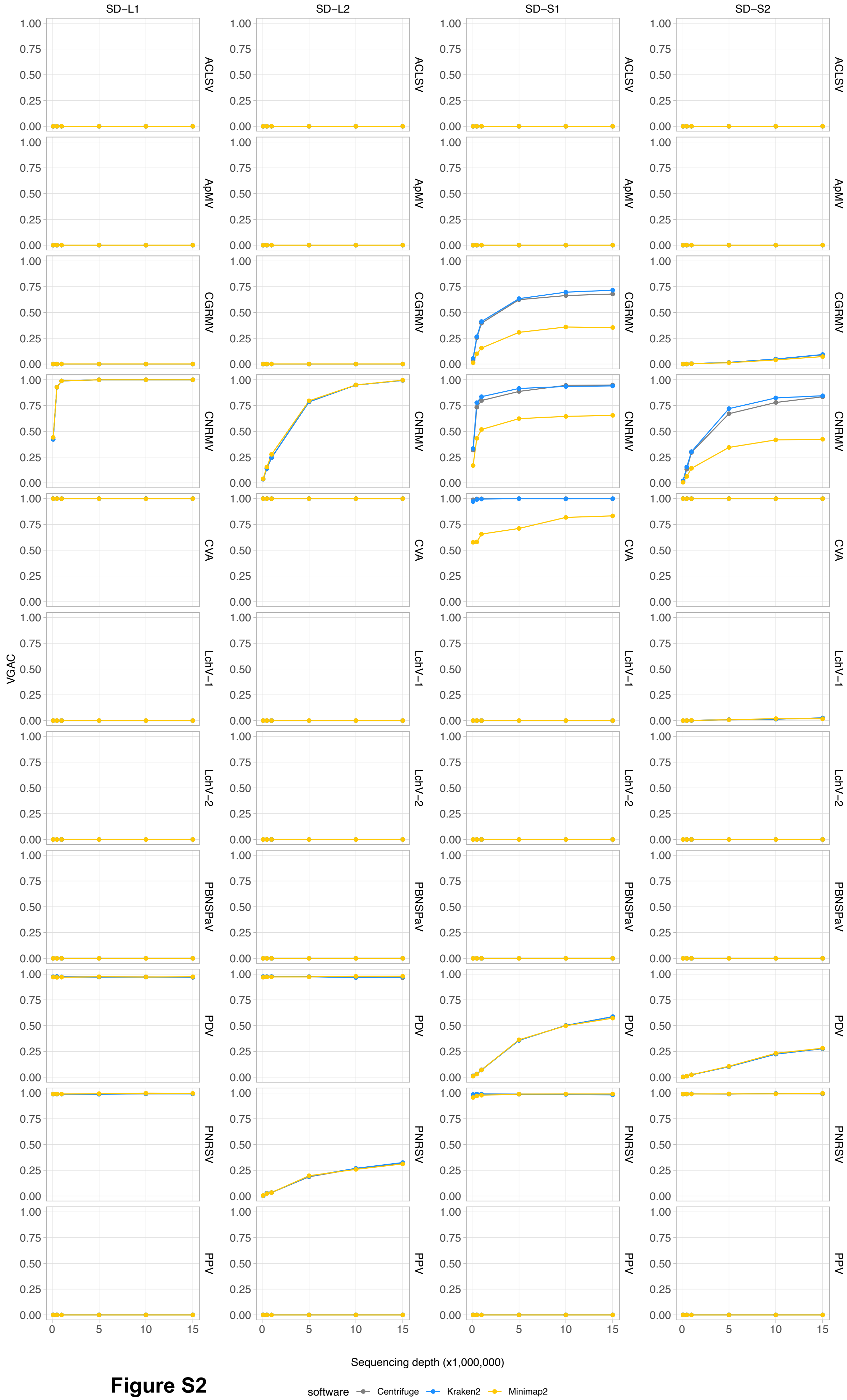
SD–L1
SD–L2
SD–S1
SD–S2
ACLSV
ApMV
CGRMV
CNRMV
CVA
LchV–1
LchV–2
PBNSPaV
PDV
PNRSV
PPV
VGAC
Sequencing depth (x1,000,000)
software
Centrifuge
Kraken2
Minimap2

**Figure S2**

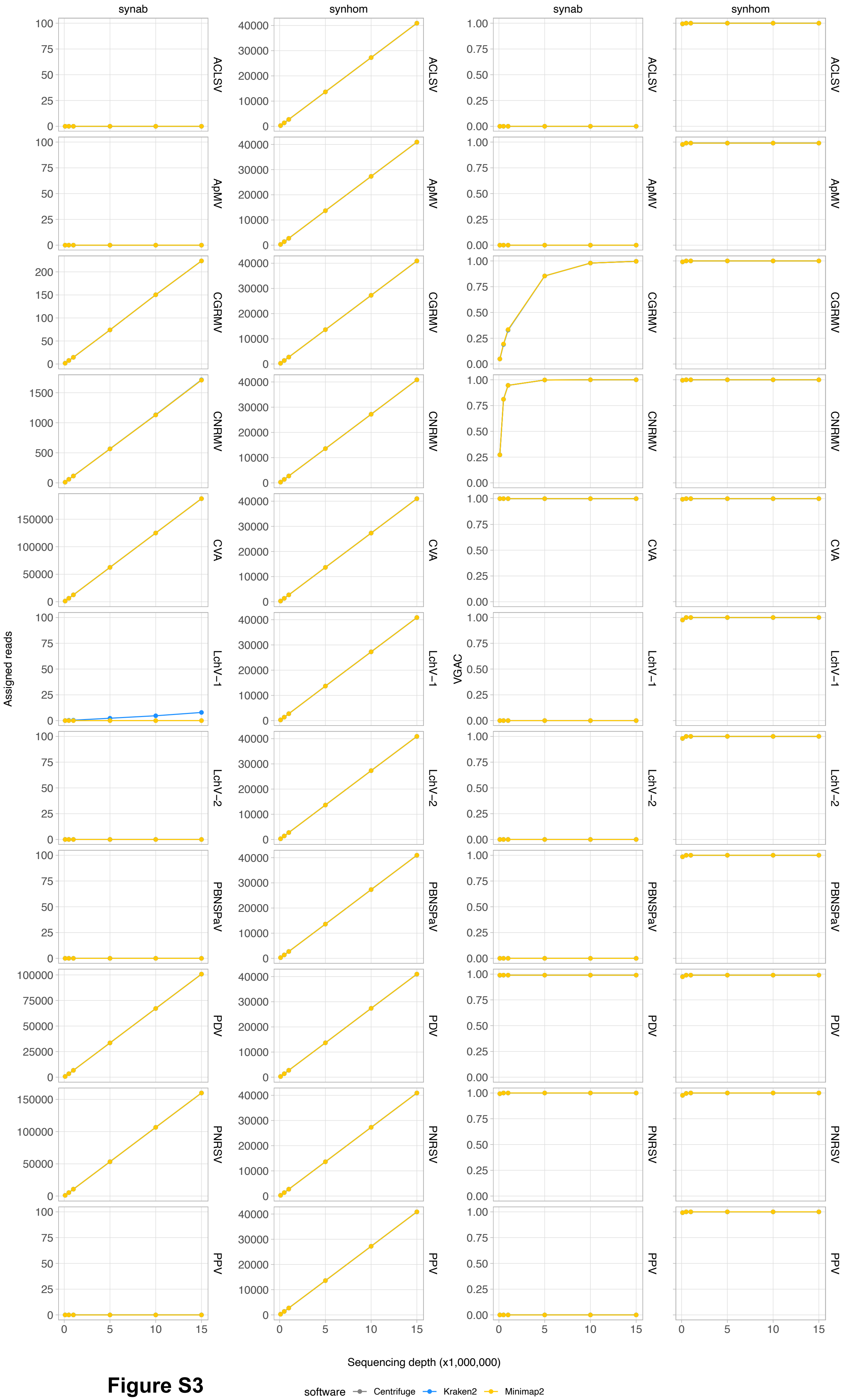
synab
synhom
synab
synhom
ACLSV
ApMV
CGRMV
CNRMV
CVA
LChV−1
LChV−2
PBNSPaV
PDV
PNRSV
PPV
Assigned reads
VGAC
Sequencing depth (x1,000,000)
software
Centrifuge
Kraken2
Minimap2

**Figure S3**

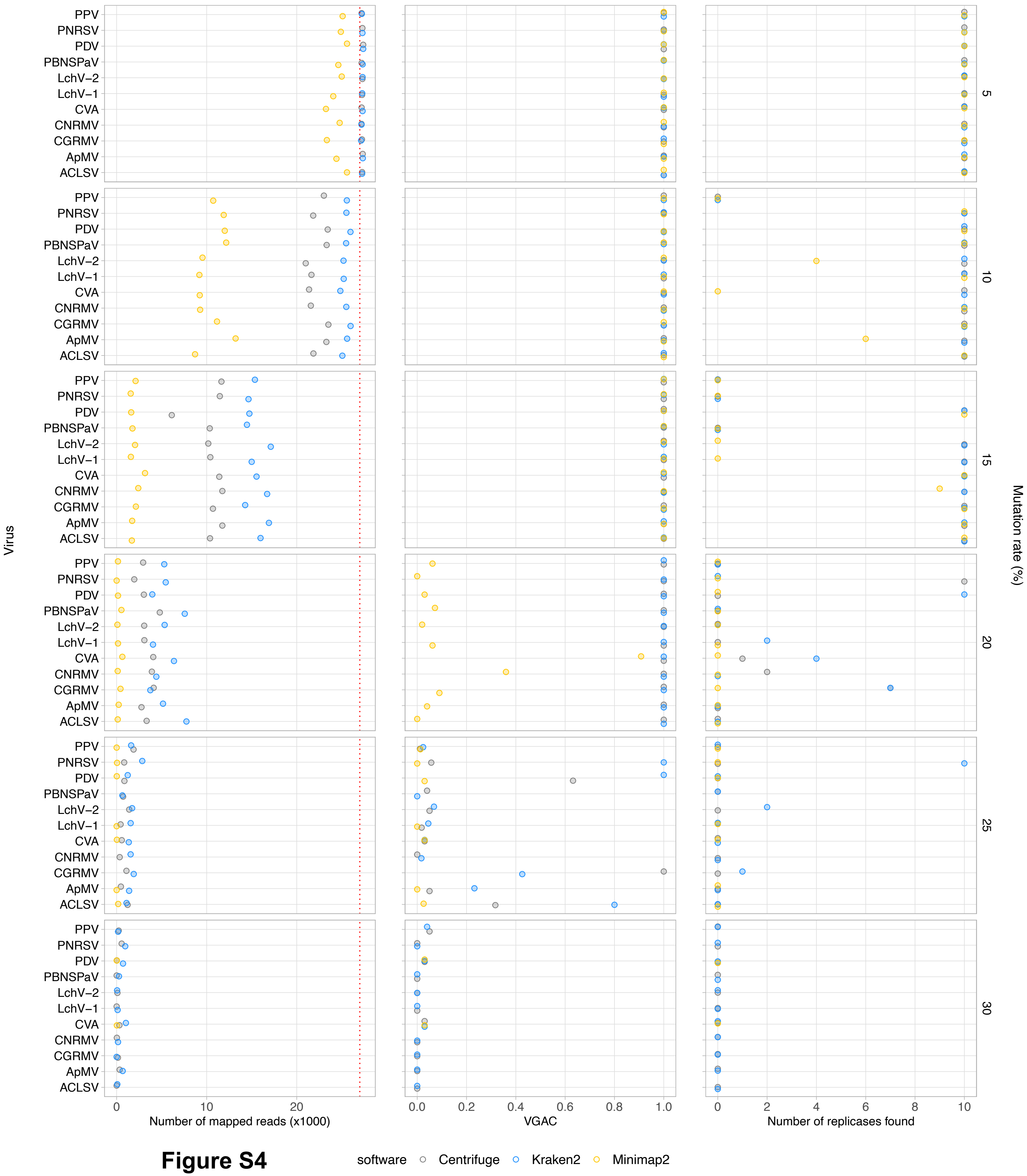
PPV
PNRSV
PDV
PBNSPaV
LchV−2
LchV−1
CVA
CNRMV
CGRMV
ApMV
ACLSV
Virus
5
10
15
20
25
30
Mutation rate (%)
0
10
20
Number of mapped reads (x1000)
0.0
0.2
0.4
0.6
0.8
1.0
VGAC
0
2
4
6
8
10
Number of replicases found
software
Centrifuge
Kraken2
Minimap2

### Figure S4

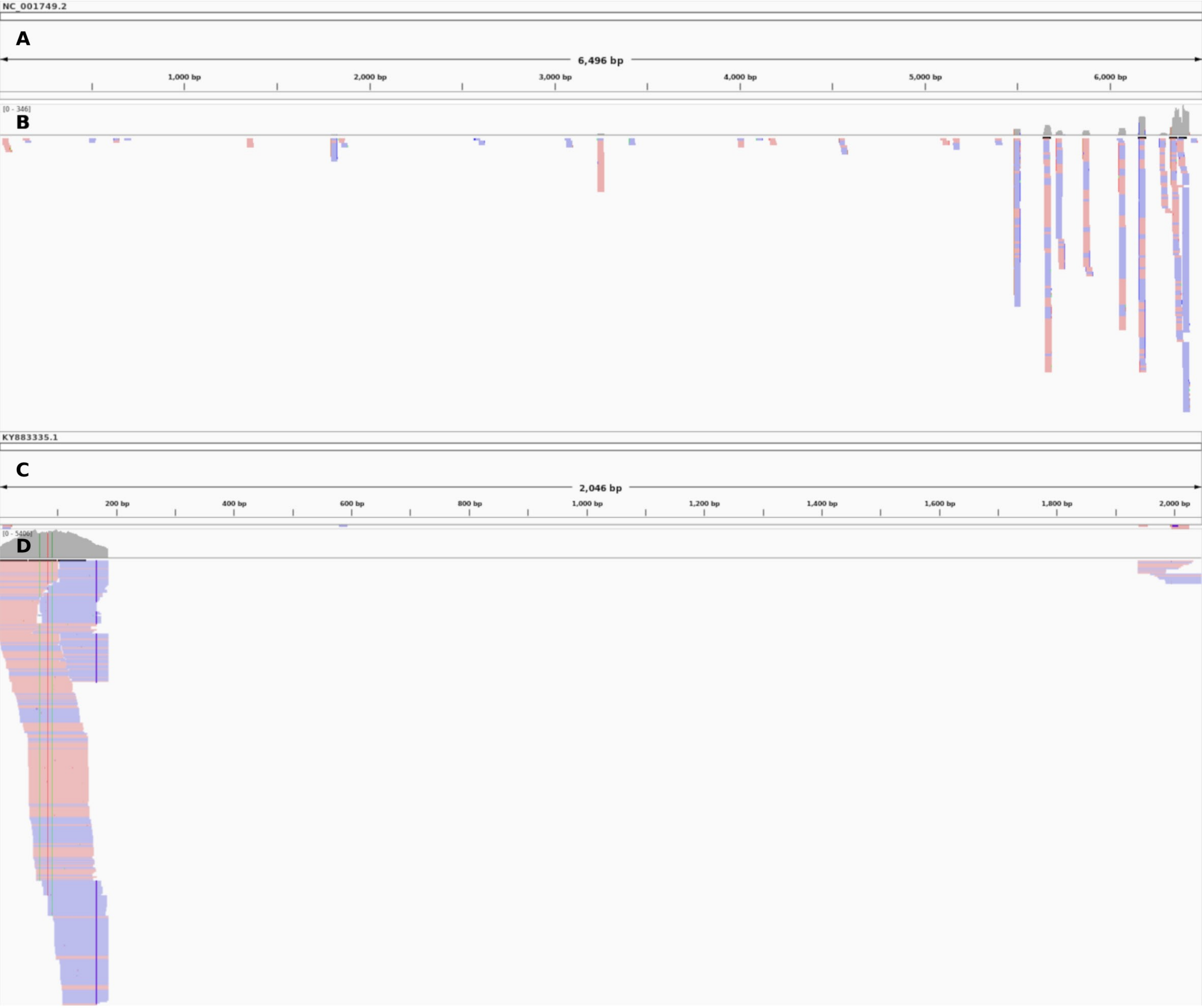

NC_001749.2
A
6,496 bp
1,000 bp
2,000 bp
3,000 bp
4,000 bp
5,000 bp
6,000 bp
[0 - 346]
B
KY883335.1
C
2,046 bp
200 bp
400 bp
600 bp
800 bp
1,000 bp
1,200 bp
1,400 bp
1,600 bp
1,800 bp
2,000 bp
D


**Figure S5**

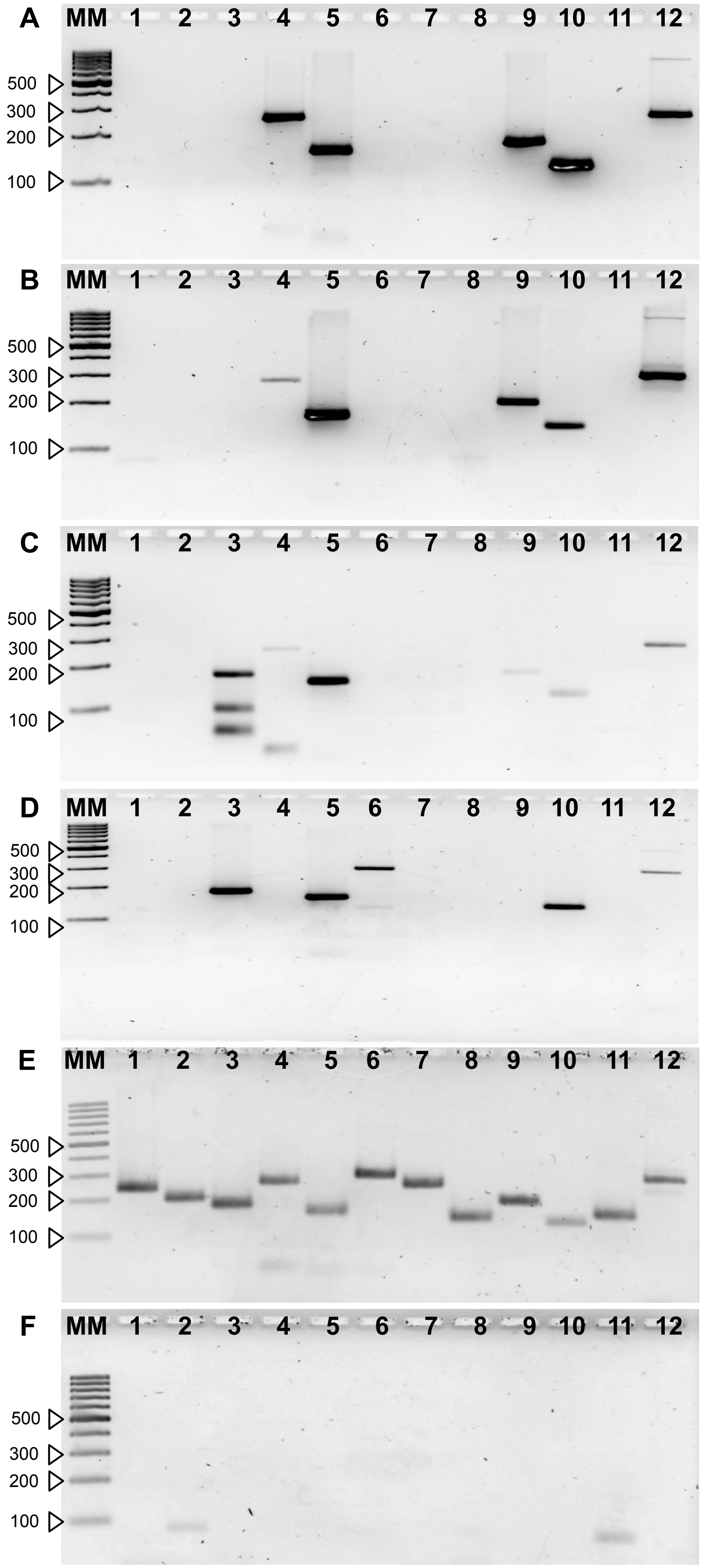
A MM 1 2 3 4 5 6 7 8 9 10 11 12
500
300
200
100
B MM 1 2 3 4 5 6 7 8 9 10 11 12
500
300
200
100
C MM 1 2 3 4 5 6 7 8 9 10 11 12
500
300
200
100
D MM 1 2 3 4 5 6 7 8 9 10 11 12
500
300
200
100
E MM 1 2 3 4 5 6 7 8 9 10 11 12
500
300
200
100
F MM 1 2 3 4 5 6 7 8 9 10 11 12
500
300
200
100

**Figure S6**

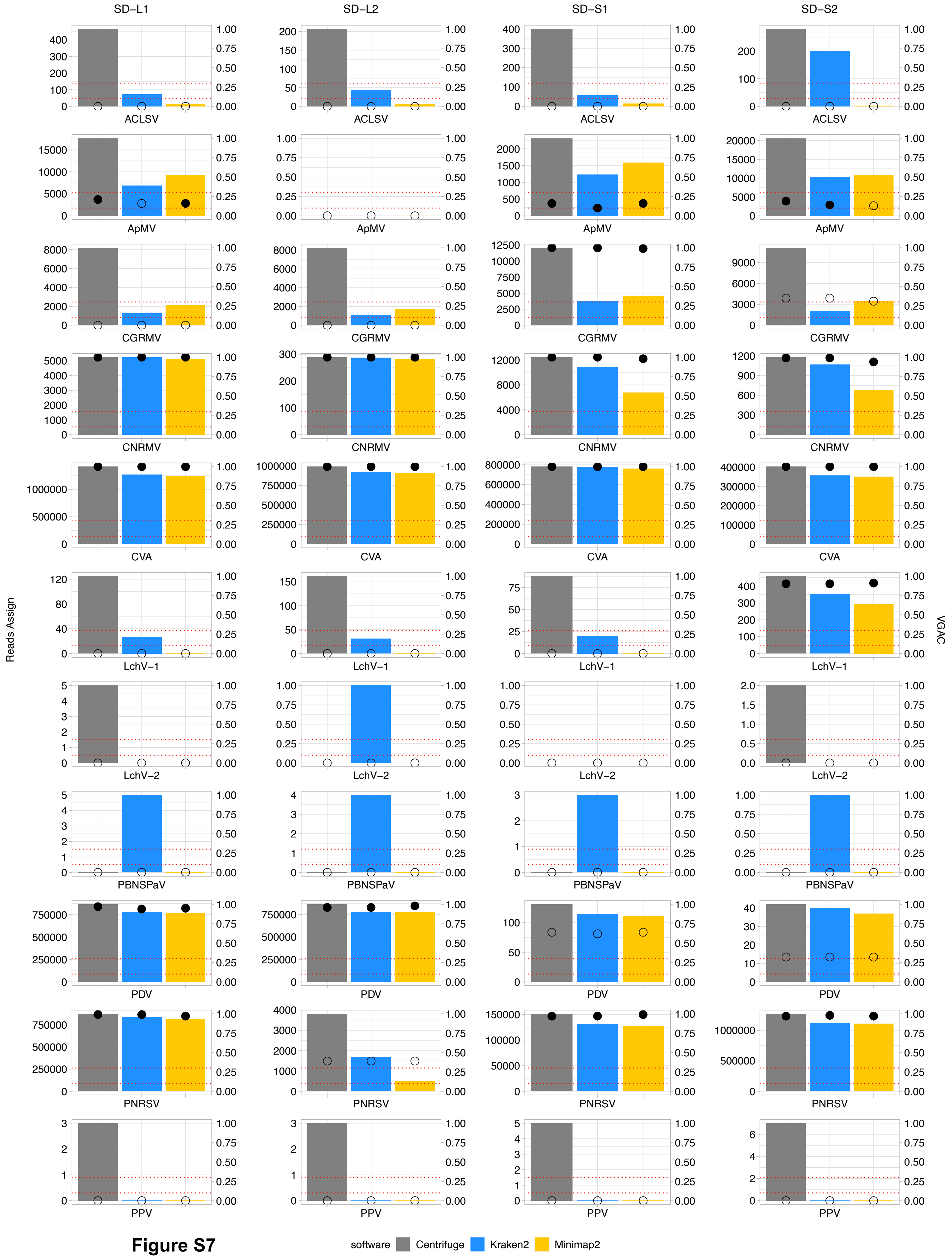
SD−L1
SD−L2
SD−S1
SD−S2
Reads Assign
VGAC
ACLSV
ApMV
CGRMV
CNRMV
CVA
LchV−1
LchV−2
PBNSPaV
PDV
PNRSV
PPV
software
Centrifuge
Kraken2
Minimap2

**Figure S7**

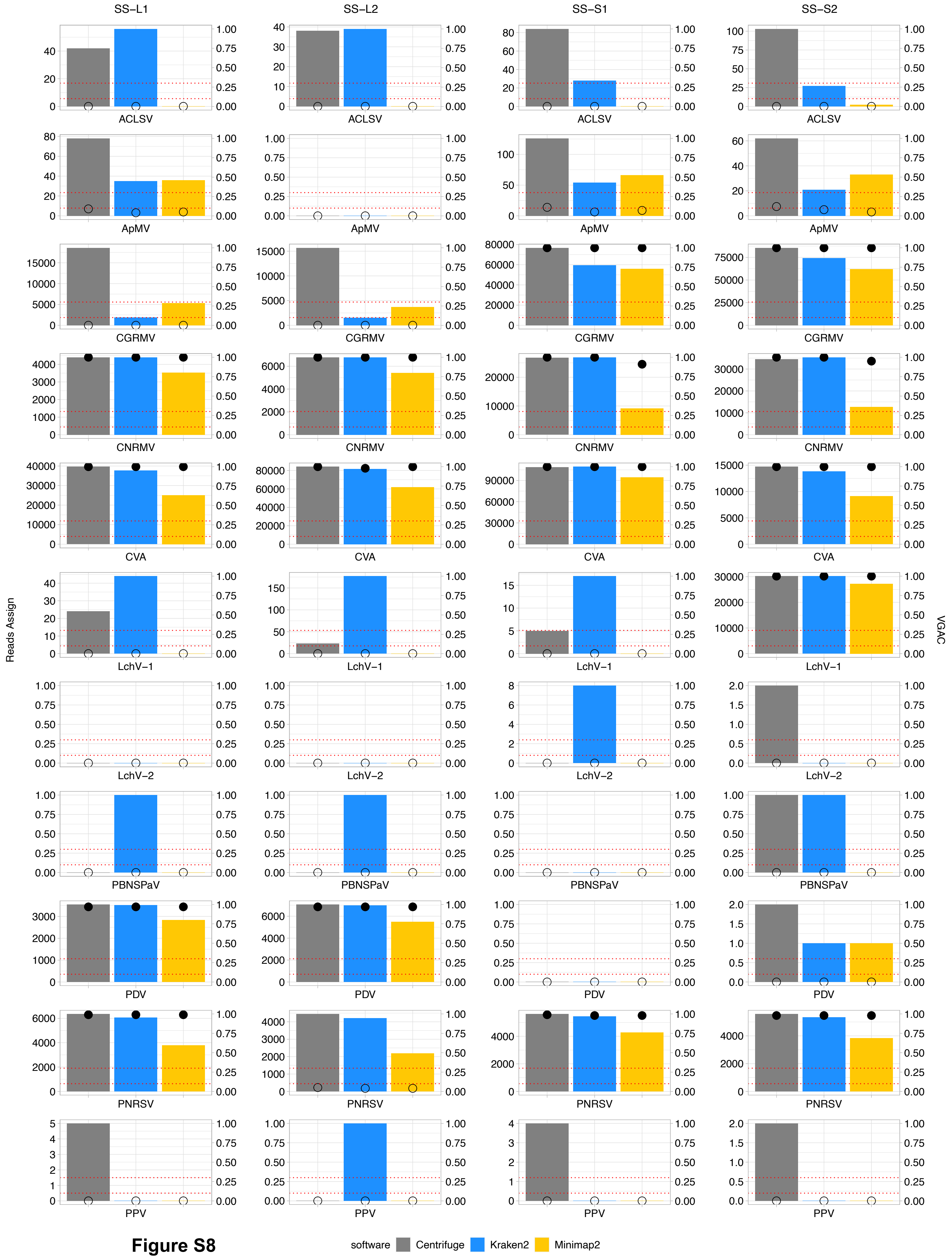
SS–L1
SS–L2
SS–S1
SS–S2
Reads Assign
VGAC
ACLSV
ApMV
CGRMV
CNRMV
CVA
LchV–1
LchV–2
PBNSPaV
PDV
PNRSV
PPV
software
Centrifuge
Kraken2
Minimap2

**Figure S8**
